# A Phosphorylation Switch Modulates Configurational Codes in the Oncofetal IGF2BP RNA Binding Paralogs

**DOI:** 10.64898/2026.05.07.723538

**Authors:** Vikas Kaushik, Vaishnavi Sanjayan, Jenna Mattice, Monika Tokmina-Lukaszewska, Ricarda Törner, Ilan Edelstein, Rahul Chadda, Rajnandani Kashyap, Abhinav Vayyeti, Pralambika Roy, Ryan A. Mehl, Richard B. Cooley, Snorri Th. Sigurdsson, Yaakov Levy, Reza Dastvan, Haribabu Arthanari, Brian B. Bothner, Sofia Origanti, Edwin Antony

## Abstract

The insulin-like growth factor 2 mRNA-binding proteins (IGF2BP1-3) are oncofetal RNA regulators that control translation, stability, and localization of several transcripts, yet display paralog-specific functions despite high structural similarity. Each paralog contains six RNA-binding domains (two RRMs and four KH domains) linked by intrinsically disordered segments. mTORC2 phosphorylates IGF2BP1 and IGF2BP3 at a single conserved serine within the disordered linker between the RRM2 and KH1 domains, a modification required for proper regulation of mRNA translational fate. Pairing site-specific phosphoserine incorporation with structural and biophysical interrogations, we show that this phosphorylation acts as a configurational switch that reorganizes long-range arrangements of RNA-binding domains and linkers without altering the secondary structure, and with only modest effects on RNA-binding affinity. Critically, pSer-driven rearrangements occur both in the RNA-free state and upon RNA engagement, and the resulting architectures differ markedly between IGF2BP1 and IGF2BP3 despite >70% sequence identity. These paralog-specific, phosphorylation-dependent configurational landscapes likely underlie differences in mRNA recognition modes and functional outcomes. Our work identifies a post-translational mechanism that tunes IGF2BP paralog dynamics across free and RNA-bound states to program target mRNA selection, processing, and translational fate.

## Introduction

mRNA binding proteins are well-known regulators of translation^1–4^. Many of these proteins possess multiple high-affinity RNA binding domains (RBDs). mRNA binding specificity is imparted through an amalgamation of weak sequence recognition, RNA structure, and protein-structural properties. The insulin-like growth factor 2 messenger RNA binding proteins (IGF2BPs or IMPs) are one such family of mRNA regulators and comprise three well-conserved paralogs - IMP1, IMP2, and IMP3^5^. IMPs function in a host of cellular processes including cell migration, metabolism, neuronal development, and stem cell renewal^6,7^. They bind to hundreds of mRNA targets each and thus have well established roles in mRNA transport & translation^8–12^, stability^13–16^, alternative splicing^17^, and m^6^A RNA methylation^18,19^. All three IMPs are abundant during embryogenesis and are important for development^20–22^. IMP1 and IMP3 expression is shut off after birth and thus largely absent in adult tissues^23,24^. IMPs are re-expressed in several cancers and correlate with poor patient survival outcomes; thus, they are defined as oncofetal in nature^25–31^. The oncofetal characteristics have rendered IMPs as attractive targets for chemotherapy. While IMPs lack catalytic activity, they serve as a scaffold for the recruitment of other proteins onto the appropriate mRNA target(s)^6,32,33^.

IMP1, IMP2, and IMP3 share a high degree of sequence and structural identity; IMP1 and IMP3 are 73.5% identical in sequence (**Figure 1**). This increases to 79% identity if the disordered linkers are excluded. The structural similarity in the IMP family raises a fundamental question about mRNA binding specificity: Specifically, how do three paralogs that share a high degree of structural similarity recognize and regulate unique mRNA targets? Spatial and temporal separation of the mRNA and the target IMP could control access to binding^34,35^. Structural features intrinsic to the mRNA and some degree of RNA-sequence preference for each RBD in IMPs have been experimentally determined^36^. Oligomerization of IMP proteins on mRNA and interaction with other proteins also regulate binding specificity^37^.

**Figure 1.**
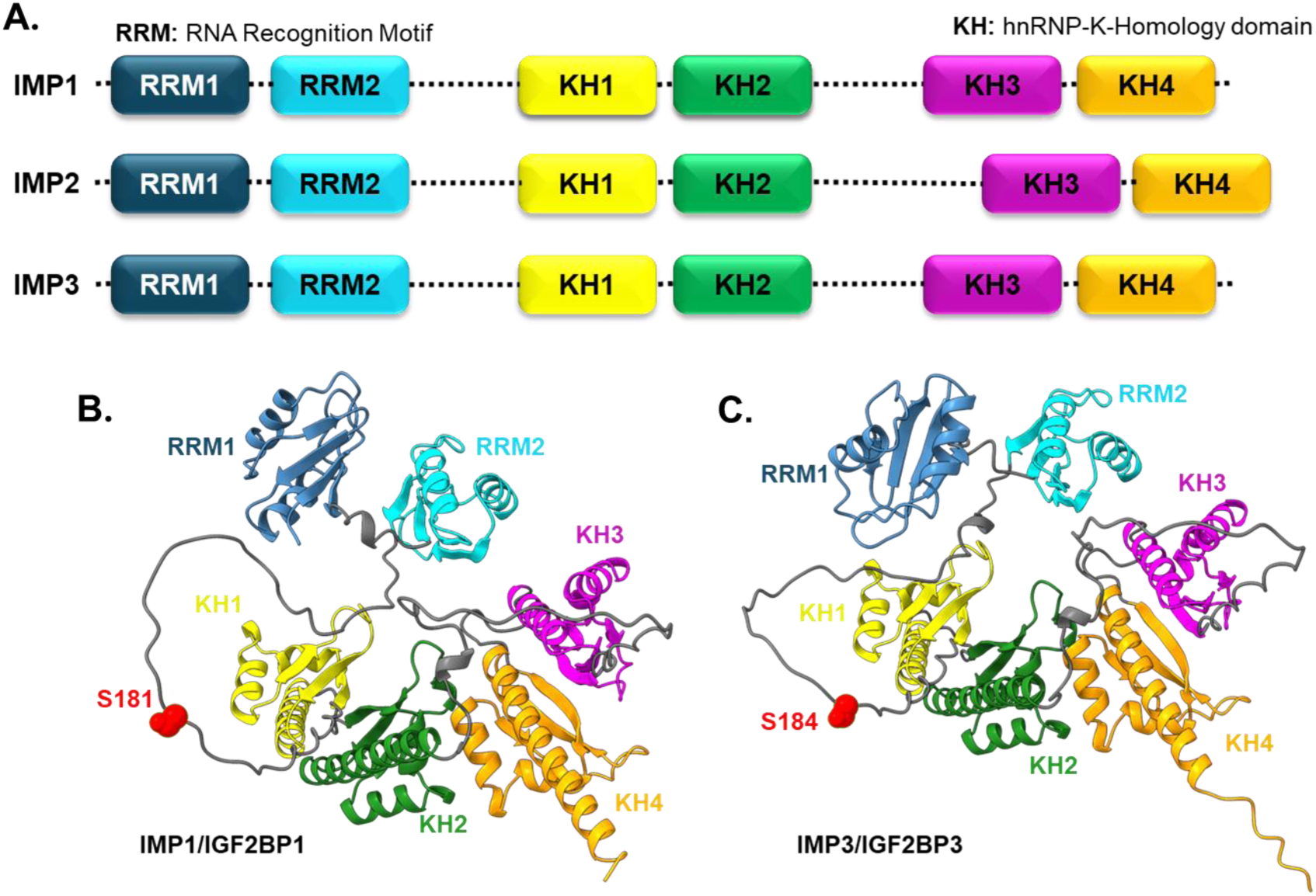
IMP paralogs share a high degree of sequence and structural similarity. **A)** All three IMP paralogs are composed of highly conserved RNA-recognition motifs (RRM1 and RRM2) and hnRNP-K-homology domains (KH1, KH2, KH3, and KH4). The dotted lines represent disordered linkers that connect the domains. AlphaFold models of **B)** IMP1 and **C)** IMP3 are shown. The individual RRM and KH-domains are depicted, and the intervening linkers are shown in grey. Ser-181 in IMP1 and Ser-184 in IMP3 are positions of phosphorylation by mTORC2 kinase and are highlighted in red.

From a structural perspective, IMPs are composed of six RBDs: two RNA recognition motif-containing domains (RRM1 and RRM2) and four hnRNP-K-homology domains (KH1, KH2, KH3, and KH4; **Figure 1**)^36^. The domains are connected by flexible linkers and those linking the RRM2-KH1 and KH2-KH3 domains are particularly long (>40 aa each). This flexible architecture allows the individual domains to adopt a wide array of configurations. We here define configurations or configurational changes as the relative position of one domain with respect to another. We refer to structural changes within a single domain as conformational changes. Configurational changes are likely regulated through physical interactions between the domains and/or the linkers. From a functional context, these configurations are also likely modulated by RNA structure and sequence accessibility and further regulated through post-translational modifications of the domains and/or the linkers.

Investigating structure-function relationships in multidomain nucleic acid binding proteins, such as IMPs, is technically challenging. For example, each of the four KH domains binds RNA with relatively low affinity when examined in isolation, yet together they mediate high-affinity RNA interactions through collective action^33^. Consequently, mutations within a single KH domain may not result in a measurable decrease in RNA-binding affinity when assessed using ensemble steady-state assays. In addition, individual KH domains exhibit weak intrinsic RNA sequence specificity^38^. Instead, the four KH domains function as pseudo-dimers, collectively recognizing target RNAs via short 4-nucleotide interaction motifs that impart specificity^6,38,39^. By contrast, the two RRM domains bind RNA more promiscuously and display even weaker sequence recognition, typically interacting with 2-nucleotide motifs^6,33,39^.

Structural studies of full-length IMP proteins are further complicated by the presence of long, flexible linker regions connecting these domains. As a result, most detailed structural and mechanistic investigations have relied on isolating and characterizing individual domains or domain pairs. Tandem domain units such as RRM1–RRM2, KH1–KH2, and KH3–KH4 have been examined using X-ray crystallography and nuclear magnetic resonance (NMR) spectroscopy and are thought to function cooperatively to recognize short nucleotide stretches, typically 3 - 6 nucleotides per unit^6,33,38–42^. However, the RNA sequence motifs identified as determinants of mRNA recognition vary substantially depending on the experimental approach employed. To date, there remains little consensus regarding the existence of ideal or highly specific sequence motifs for the IMP paralogs^33,36,39,43,44^. These observations suggest that RNA recognition by IMPs is likely governed by cooperative interactions among multiple domains, in conjunction with features of RNA secondary or higher-order structure. While studies of individual domains have yielded valuable insights, they do not account for the direct and allosteric communication across domains observed in the full-length protein, leading to incomplete models of RNA-binding specificity and mechanism. More recent studies have therefore shifted toward the use of advanced biophysical and structural techniques to interrogate functional specificity in the context of full-length IMP-RNA complexes^33,45^.

Finally, post-translational modifications of IMPs have been identified and proposed to further refine mRNA binding^45–47^. One well-established regulatory mechanism for IMP1 and IMP3 is phosphorylation by the mTOR complex 2 (mTORC2)^48^. During embryogenesis, IMP1 and IMP3 bind both the 5′ and 3′ untranslated regions (UTRs) of insulin-like growth factor 2 (IGF2) mRNA. Both proteins are phosphorylated at a single site within the disordered linker connecting RRM2 and KH1 (**Figures 1B and C**): Ser-181 in IMP1 and Ser-184 in IMP3. Loss of this phosphorylation in IMP1 disrupts proper splicing of IGF2 mRNA, leading to reduced translation and impaired cell proliferation^48^. The phosphorylation state of IMP1 has also been shown to dictate its interactions with specific mRNA targets. For example, mTORC2-mediated phosphorylation at Ser-181 promotes IMP1 binding to the 3′ UTR of ACTB mRNA and the 5′ UTR of IGF2 mRNA^49^. Notably, these interactions have opposing translational outcomes: ACTB translation is inhibited, whereas IGF2 translation is enhanced^47^. In contrast, phosphorylation of IMP1 at Tyr-396 by Src kinase promotes release from ACTB mRNA, resulting in increased ACTB translation^5,47^. Additionally, IMP1 phosphorylation influences its propensity for RNA-dependent condensate formation and contributes to its roles within ribonucleoprotein granules and P-bodies^45^. These findings suggest a complex interplay between phosphorylation of IMP proteins to enact distinct translational outcomes of specific mRNA targets^49–51^. Such functional investigations for Ser-184 phosphorylation in IMP3 have not been performed. However, the mechanisms that govern this specificity are poorly understood.

Here, using full-length IMP1 and IMP3, as well as genetic code expansion-based site-specific incorporation of phosphoserine (pSer), we investigate how phosphorylation within a disordered loop alters mRNA binding and recognition. Specifically, we probe whether phosphorylation at similar positions in two proteins with very high structural and sequence identity generates comparable structural alterations. We find that the configurational arrangements of the RNA-binding domains (RBDs) are significantly reordered upon phosphorylation and differ between IMP1 and IMP3. Our findings reveal how post-translational modifications contribute to mRNA recognition and may differentially alter functional specificity in IMP1 and IMP3.

## Results

### Phosphorylation at the mTORC2 sites moderately affects the RNA binding affinity of both IMP1 and IMP3

mTORC2 phosphorylates IMP1 and IMP3 at Ser-181 or Ser-184, respectively. These positions reside in the disordered linker connecting the highly conserved RRM2 and KH1 domains (**Figure 1B & C**). To study the effect of mTORC2 phosphorylation on the structural and biochemical properties of IMP1 and IMP3, we utilized genetic code expansion to site-specifically incorporate phosphoserine (pSer)^52,53^. This efficient method produces >95% pSer carrying IMP1 and IMP3 as evidenced by the slower migration of the pSer-carrying proteins on a Phos-Tag SDS PAGE gel (**Figures 2A & B**). Site-specific incorporation of pSer at the designated positions in IMP1-pSer^181^ and IMP3-pSer^184^ was further confirmed by mass spectrometry (**Figures 2C & D**). These data show stable and complete pSer incorporation in both proteins and thus facilitate the structural and biophysical studies described below.

**Figure 2.**
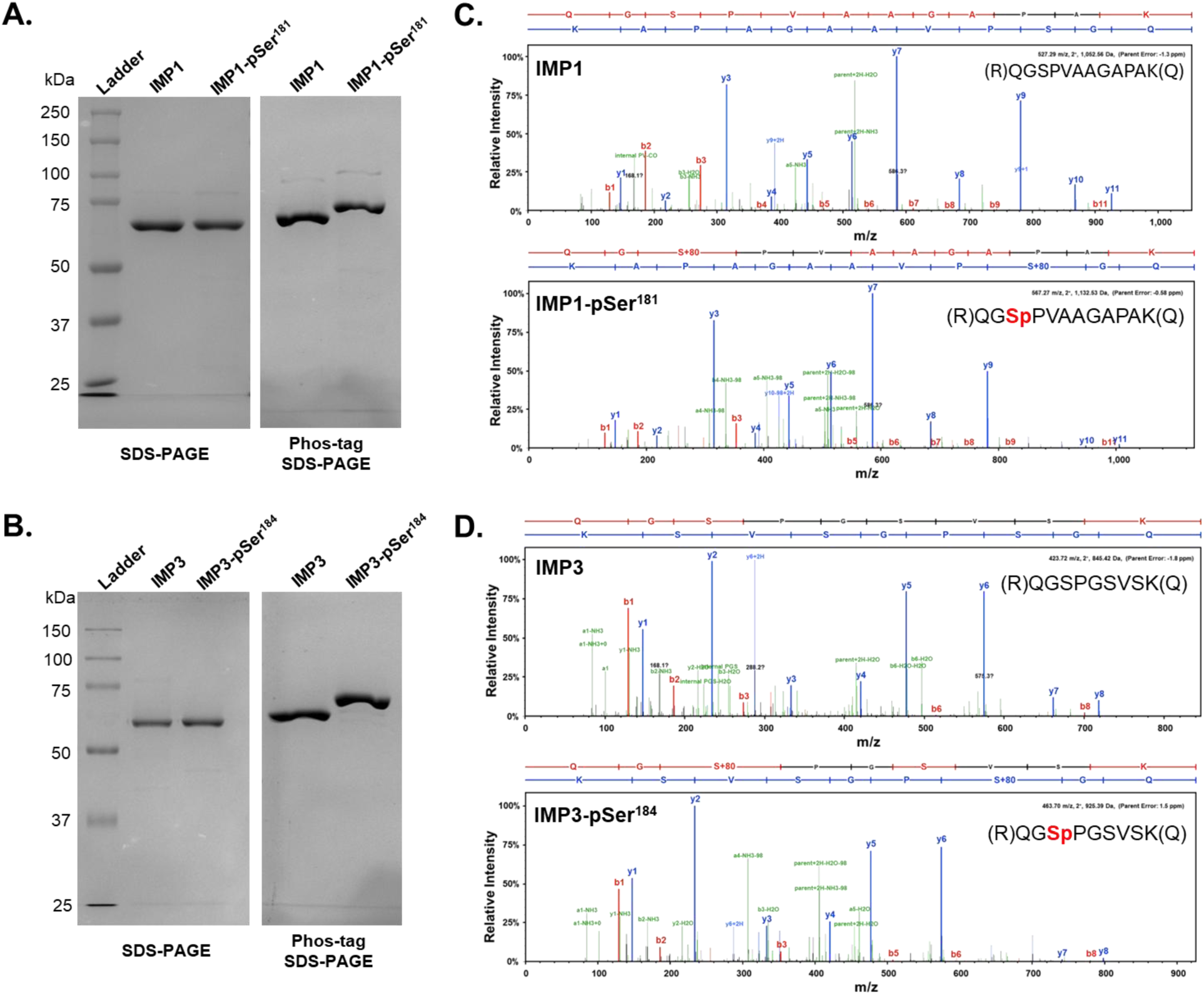
Phosphoserine incorporation in IMP1 and IMP3. SDS-PAGE and Phos-tag SDS PAGE analysis of recombinant **A)** IMP1 and **B)** IMP3. Both the unmodified and pSer-modified proteins migrate similarly on normal SDS-PAGE, but both IMP1-pSer^181^ and IMP3-pSer^184^ show slower electrophoretic mobility in Phos-tag SDS PAGE analysis. Mass spectra confirm site-specific pSer incorporation at positions **C)** 181 in IMP1 and **D)** 184 in IMP3.

Fluorescence anisotropy analysis was used to investigate the effect of phosphorylation on RNA binding affinity of IMP1 and IMP3. RNA substrates carrying 5′ fluorescein were used and the change in anisotropy was measured as a function of protein concentration. For IMP1, we used a 54 nt RNA sequence from a *cis*-acting element that immediately follows the termination codon in the 3′ untranslated region (UTR) of β-actin mRNA (Zipcode)^42,54^. For IMP3, we used a longer 121 nt substrate from a region in exon 29 of ANKRD17 mRNA (ANKRD)^33,55^. Both RNA substrates are biochemically well-characterized targets for IMP1 and IMP3 binding^6,33,55,56^. IMP1 and IMP3 bind to the cognate RNA substrate with high affinity (**Figures 3A & B**). IMP1-pSer^181^ binds with marginally lower affinity (K_d_ = 17.3±3.1 nM) compared to IMP1 (K_d_ = 6.7±1.7 nM). Similarly, IMP3-pSer^184^ binds its cognate RNA substrate with slightly lower affinity (K_d_ = 58.4±15.5 nM) compared to IMP3 (K_d_ = 17.2±4.2 nM). However, it should be noted that all these interactions are in the low nM range and therefore should be classified as high-affinity interactions.

**Figure 3.**
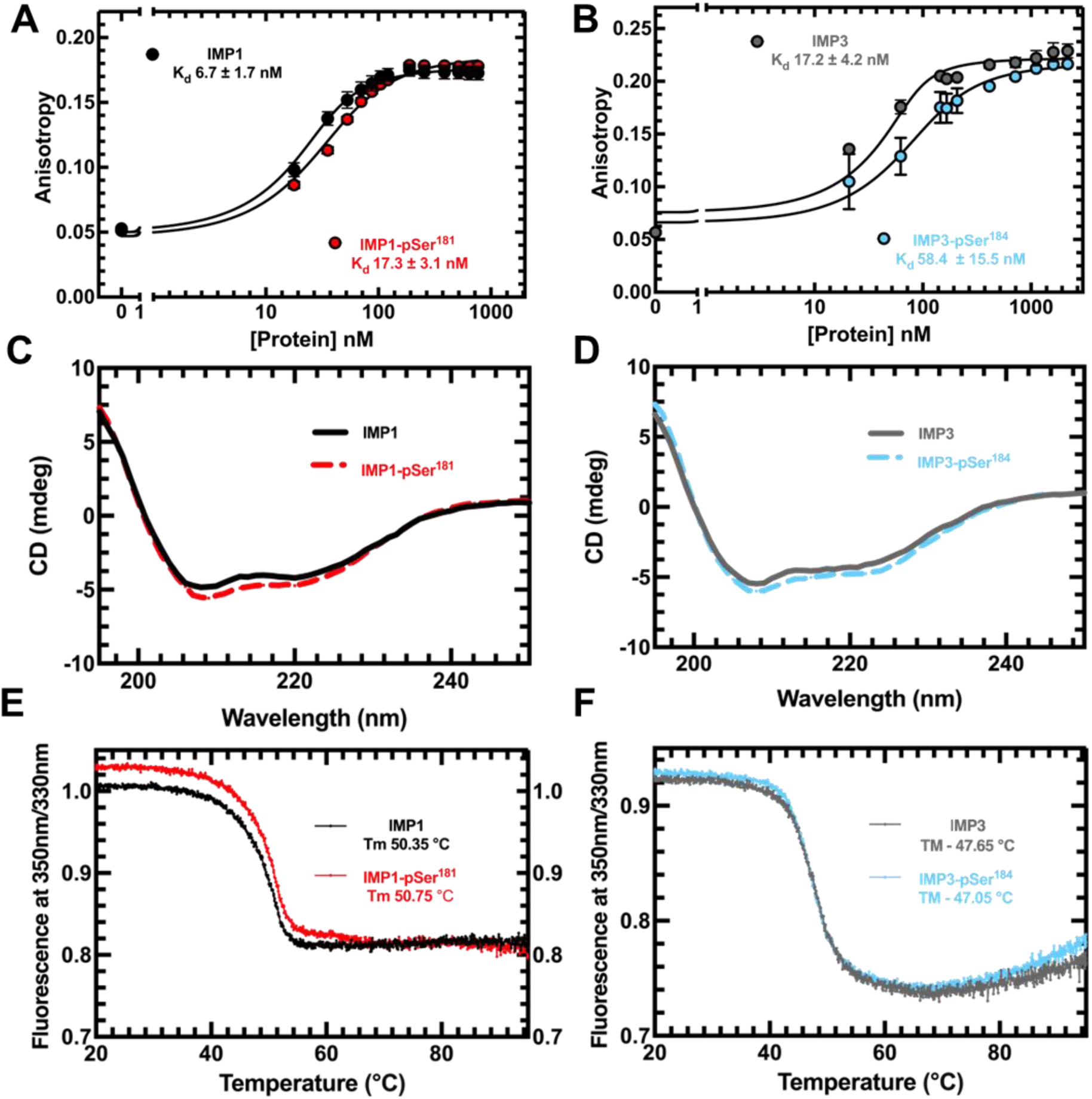
Phosphorylation has marginal effects on the RNA binding and secondary structure properties of IMP1 and IMP3. Fluorescence anisotropy analysis of RNA binding to **A)** IMP1 or IMP-pSer^181^ and **B)** IMP3 or IMP3-pSer^184^ show moderate reduction in RNA binding affinity upon phosphorylation. Data from three independent experiments are shown and error bars denote SEM. Circular dichroism spectra of **C)** IMP1 and IMP1-pSer^181^ and **D)** IMP3 or IMP3-pSer^184^ show no appreciable changes in the secondary structures upon phosphorylation. Nano-differential scanning fluorimetry (nanoDSF) analysis of **E)** IMP1 or IMP1-pSer^181^ and **F)** IMP3 or IMP3-pSer^184^ show no appreciable differences in the melting temperature suggesting no quantifiable changes in the secondary structure upon phosphorylation. Representative CD and nanoDSF data from three independent experiments are shown.

### Phosphorylation does not introduce large-scale changes in the secondary structure of the RNA-binding domains of IMP1 or IMP3

Ser-181 in IMP1 and Ser-184 in IMP3 reside in the disordered linker connecting the RRM2 and KH1 domains (**Figures 1B & C**). Interactions between the linkers and the folded RRM and KH domains have been reported^45^. Therefore, we tested whether introduction of pSer altered the overall secondary structure of the proteins using circular dichroism (CD). The overall CD spectral profiles of IMP1 and IMP1-pSer^181^, and that of IMP3 and IMP3-pSer^184^ are similar (**Figures 3C & D**). Analysis of the CD profiles show a ∼10% increase in the helical content for IMP1-pSer^181^ over IMP1 (**Supplemental Table 3**). In contrast, we observe a ∼10% decrease in helical content for IMP3-pSer^184^ over IMP3 (**Supplemental Table 3**). Furthermore, nano-differential scanning fluorimetry analysis of the proteins show no difference in the melting temperatures for IMP1 and IMP1-pSer^181^ or IMP3 and IMP3-pSer^184^, respectively (**Figures 3E & F**). These data suggest that pSer introduction at these positions does not generate large-scale changes in the overall secondary structure of the well-conserved RNA binding domains in either IMP1 or IMP3. However, the CD data do suggest that upon phosphorylation, the linkers might be generating configurational changes that are unique to IMP1 versus IMP3.

To get a higher resolution picture of any structural changes induced by phosphorylation, we probed IMP1 and IMP1-pSer^181^ using NMR spectroscopy. The ^15^N-TROSY-HSQC spectrum is characteristic of a multidomain protein comprising folded domains connected by intrinsically disordered linkers. Intense, poorly dispersed resonances in the random-coil region of the ^1^H dimension, approximately 7.5–8.5 ppm, are consistent with the intrinsically disordered linkers, whereas weaker upfield- and downfield-shifted resonances arise from the folded domains (**Figure 4A**). An overlay of the ^15^N-TROSY-HSQC spectra of IMP1 and IMP1-pSer^181^ shows that selective and complete phosphorylation of position 181 was achieved (**Figure 4A**). A large downfield chemical shift perturbation (CSP) was observed for a single resonance in the spectral region typically populated by serine and threonine resonances from intrinsically disordered polypeptide segments. This resonance shifted from 117.5 ppm in ^15^N and 8.1 ppm in ^1^H to 119.6 ppm in ^15^N and 8.6 ppm in ^1^H. Based on this characteristic behavior, the residue was assigned to S181 (**Figure 4A**). Only a few chemical shift perturbations were observed above 8.5 ppm or below 7.5 ppm in the ^1^H dimension, where resonances corresponding to structured regions are typically found. Some resonances in these regions exhibited broadening upon phosphorylation, likely reflecting chemical exchange. This behavior is consistent with phosphorylation-dependent changes in how the intrinsically disordered linker interacts with the folded domains. A few resonances in the region corresponding to disordered polypeptide segments, (^1^H 7.5–8.5 ppm), also displayed CSPs upon phosphorylation. These CSPs may arise from residues neighboring the phosphorylated site or from residues involved in long-range interactions with the phosphorylated residue, as indicated by arrows in **Figure 4A**. Taken together, the NMR observations suggest that S181 phosphorylation does not substantially perturb the global fold of the structured regions of IMP1, consistent with the orthogonal CD and nanoDSF measurements (**Figures 3C & E**). However, we cannot exclude the possibility that phosphorylation induces local conformational changes within the disordered region. The observation of resonances from the folded regions of a 65 kDa protein in the absence of deuteration suggests that the folded domains undergo at least partially independent tumbling.

**Figure 4.**
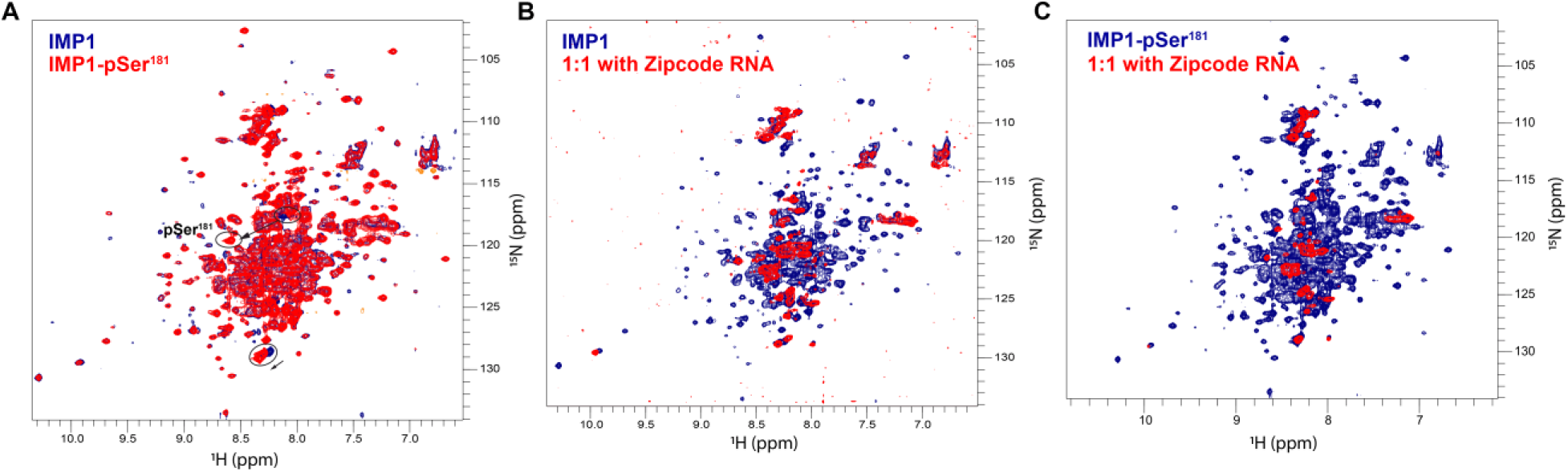
¹H–¹⁵N TROSY-HSQC spectral overlays of IMP1, IMP1-pS181, and their complexes with Zipcode RNA. **A)** Overlay of unmodified IMP1 (blue) and Ser181-phosphorylated IMP1, IMP1-pSer^181^ (red). **B)** Overlay of IMP1 alone (blue) and IMP1 in the presence of Zipcode RNA at a 1:1 molar ratio (red). **C)** Overlay of IMP1-pSer^181^ alone (blue) and IMP1-pSer^181^ in the presence of Zipcode RNA at a 1:1 molar ratio (red). S181 phosphorylation results in only limited chemical shift perturbations, consistent with no major perturbation of the folded domains. In contrast, addition of Zipcode RNA causes pronounced resonance broadening and loss of signal intensity, consistent with RNA binding and formation of higher-molecular-weight complexes.

### IMP1, IMP3, and the phosphorylated versions are predominantly monomers in solution, but dimerize on RNA

Determination of the oligomerization properties of IMP1, IMP3 and their pSer-versions is a prerequisite to decipher the structure-function relationships. In addition, IMP1 has been shown to dimerize on RNA by the sequential binding to two protein molecules^37^. Therefore, we performed mass photometry analysis of all four proteins (IMP1, IMP-pSer^181^, IMP3, and IMP3-pSer^184^) in the absence or presence of their cognate RNA substrates (**Figure 5, Supplementary Figure 2, and Supplementary Table 4**). IMP1 is predominantly a monomer in solution with a small fraction (∼14%) of dimers (**Figure 5A**). IMP1-pSer^181^ shows the presence of monomers, a small fraction of dimers (∼19%), and a minor fraction of higher order trimers (5%) (**Figure 5B**). These data suggest that the configuration of the domains/linkers in IMP1-pSer^181^ might be more accessible to promote oligomerization. In contrast, IMP1 and IMP1-pSer^181^ dimerize on the Zipcode-RNA (**Figures 5C-E**). The predicted mass of the Zipcode RNA (17.6 kDa) is below the limit of reliable measurement by MP. These data are in agreement with previous studies showing that IMP1 does not dimerize in solution, but sequentially bind to RNA to form higher order RNA-driven dimers^37^.

**Figure 5.**
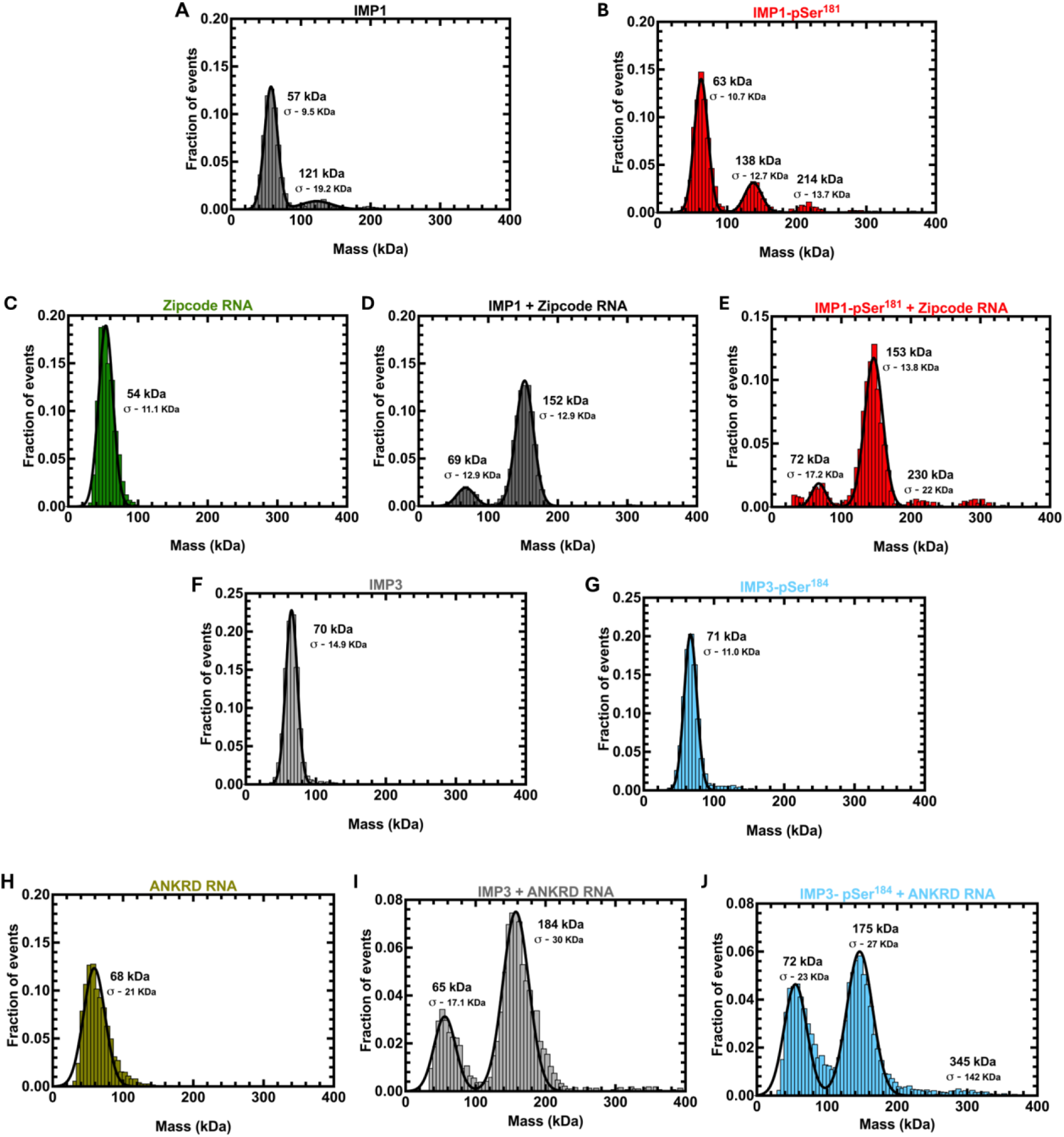
Phosphorylation induces minor changes in the oligomerization properties of IMP1 and IMP3. Mass photometry (MP) analysis of **A)** IMP1 and **B)** IMP1-pSer^181^ shows a predominantly monomeric species. Expected mass of an IMP1 monomer is 64.3 kDa. A smaller population corresponding to an IMP1 dimer is also observed. The fraction of dimers is increased upon phosphorylation along with formation of a trimeric species. Mass photometry analysis of the **C)** Zipcode RNA, and in complex with **D)** IMP1 or **E)** IMP1-pSer^181^. The mass of the Zipcode RNA (17.6 kDa) is below the limit of accurate estimation by MP. IMP1 and IMP1-pSer^181^ form dimers on RNA. **F)** IMP3 and **G)** IMP3-pSer^184^ are predominantly monomeric. **H)** The longer ANKRD (121 nt; 38.7 kDa) forms dimers in solution and two molecules of **I)** IMP3 or **J)** IMP3-pSer^184^ are bound to the RNA.

In the NMR analysis, HN-TROSY-HSQC spectra of IMP1 recorded in the presence of equimolar Zipcode RNA showed clear evidence of binding, consistent with the high affinity measured by fluorescence anisotropy (**Figure 3A**). The most striking feature of the RNA-bound spectrum is the pronounced peak broadening in spectral regions corresponding to backbone amide resonances from folded protein regions, including the RNA-binding domains (**Figure 4B**). Upon addition of RNA, only side-chain resonances and a subset of peaks arising from disordered regions remained detectable. Given the multimerization observed upon RNA binding, these spectral changes can be rationalized by the formation of higher-molecular-weight complexes with slower tumbling rates, leading to signal broadening through enhanced relaxation. Chemical exchange governed by the on and off rates of RNA binding may also contribute to signal broadening or disappearance. However, the near-complete broadening of resonances from the folded domains is more consistent with oligomerization and formation of higher-molecular-weight complexes than with chemical exchange alone. Very similar spectral changes were observed for IMP1-pSer^181^ in the presence of Zipcode RNA (**Figure 4C**), consistent with its comparable RNA affinity and similar multimerization behavior upon binding.

IMP3 and IMP3-pSer^184^ are both strictly monomeric in the absence of RNA (**Figures 5F & G**). However, similar to IMP1, IMP3 and IMP3-pSer^184^ form dimers on the ANKRD RNA substrate (**Figures 5H-J**). Interestingly, IMP3 shows a higher propensity to form the dimeric-RNA complex (74%) compared to IMP3-pSer^184^ (53%). A diffuse population of monomeric IMP3-pSer^184^-RNA bound species is also observed that is absent for the non-phosphorylated form. These properties are also observed at higher ratios of IMP3/ IMP3-pSer^184^ over RNA (**Supplementary Figure 2 and Supplementary Table 4**). These data suggest that the pSer modified IMP1/IMP3 proteins likely form alternate configurations on and off the RNA whilst retaining the secondary structure conformations of the folded RBDs.

### Distinct large-scale domain and linker rearrangements are introduced upon phosphorylation of IMP1 and IMP3

To obtain a coarse-grain structural assessment of the configurational changes, we performed cross-linking coupled mass spectrometry (XL-MS) using bis(sulfosuccinimidyl)suberate (BS3). The *N*-hydroxysulfosuccinimide (NHS) esters positioned at either end of BS3 are spaced 8-atoms apart and report on crosslinks with primary amines that are within ∼12 Å^57,58^. For IMP1, several crosslinks (XLs) are captured between the four KH domains. In the IMP1-pSer^181^ protein, many of the XLs observed in IMP1 are lost. Interestingly, all the crosslinks originating from the intrinsically disordered linkers around the site of phosphorylation are lost, suggesting major configurational rearrangements. In the presence of RNA, the XL patterns differ even more. Comparison of the IMP1 and IMP1-RNA data reveal that XLs between KH1 and KH3 & KH4 are lost along with XLs between KH2 and KH3 & KH4 domains (**Figure 6**). In contrast, comparison of the IMP1-pSer^181^ and IMP1-pSer^181^-RNA data reveal that most XLs are lost upon RNA binding (**Figure 6**). The RRM1 domain appears to move closer to KH4 upon RNA binding to IMP1-pSer^181^. A comparison of the RNA-bound states of IMP1 and IMP1-pSer^181^ shows more XLs in the non-phosphorylated version. Collectively, these data show that IMP1 and IMP1-pSer^181^ adopt different configurations and these lead to differences in the structures of the RNA bound states.

**Figure 6.**
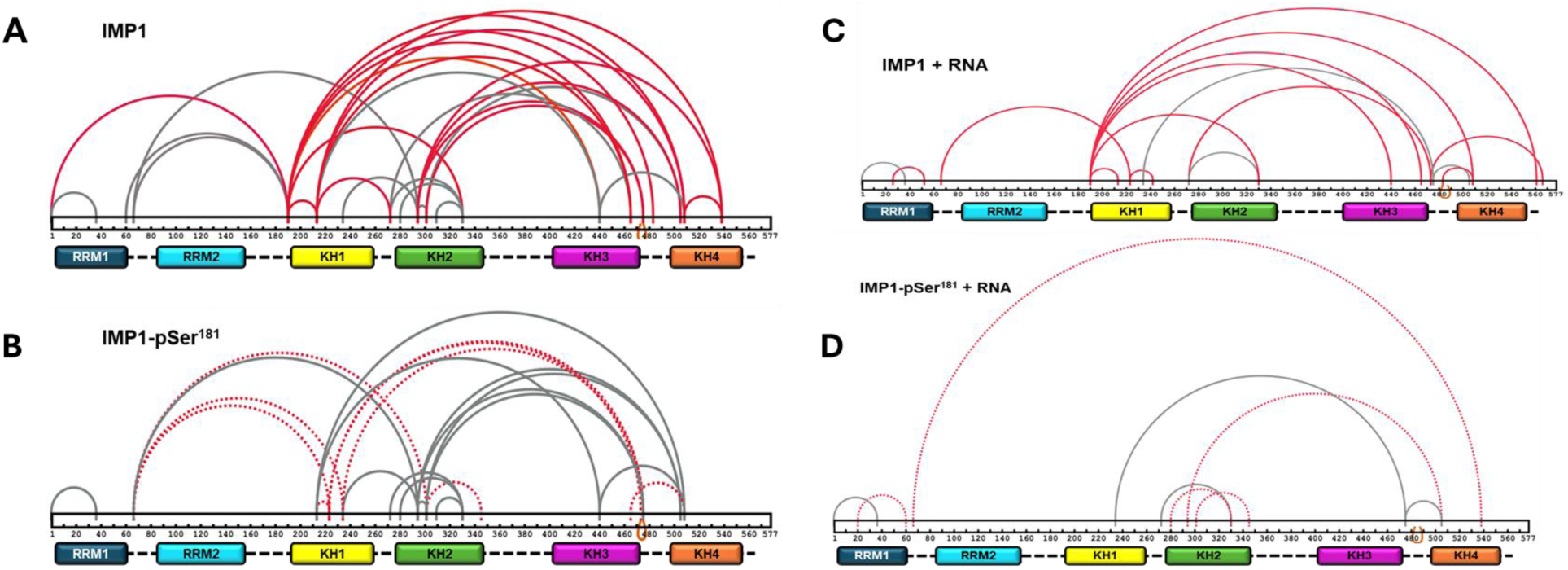
Cross-linking mass spectrometry analysis of IMP1 and IMP1-pSer^181^ reveal configurational changes induced upon phosphorylation. Crosslink maps of **A)** IMP1 and **B)** phosphorylated IMP1 (IMP1-pSer^181^) in the absence or **C-D)** presence of Zipcode RNA are shown. Cross-links identified using the amine-reactive cross-linker BS3 are displayed. Arcs denote cross-linked residues along the primary sequence, with the domain architecture of IMP1 (RRM1– 2 and KH1–4) shown below each map. The crosslinking data collected in the absence and presence of RNA were analyzed as pairs. Red arcs indicate cross-links unique to the indicated condition, whereas gray arcs represent cross-links shared between the pair in each scenario. Phosphorylation at Ser181 results in a significant redistribution of inter-domain cross-links, consistent with configurational rearrangements that are further modulated by RNA binding.

Given the >70% sequence identity between IMP1 and IMP3 we investigated whether the configurational changes driven by phosphorylation were similar by carrying out XL-MS on the IMP3 and IMP3-pSer^184^ proteins (**Figure 7**). There are some remarkable differences in the crosslinking patterns between IMP1 and IMP3 and their respective pSer versions. First, a comparison between IMP3 and IMP3-pSer^184^ shows a much substantially higher number of XLs in the phosphorylated protein (**Figure 7**). Specifically, the region around position 184 in the IDL makes contacts with KH3, KH4, and the IDL connecting the two domains. In addition, new XLs arise between RRM1 and the adjacent IDL and the KH3-KH4 domains. The enhanced pattern of XLs upon phosphorylation suggests an extensive configurational rearrangement of IMP3-pSer^184^ compared to IMP3. The patterns of XLs in IMP3 and IMP3-pSer^184^ are vastly different from IMP1 and IMP1-pSer^181^ indicating that the configurational arrangements are different although they share a high degree of sequence identity. Upon binding to RNA, both IMP3 and IMP3-pSer^184^ display an overall loss in XLs (**Figure 7**). The patterns of XLs are also different between the unbound and RNA-bound complexes. These data suggest that differences in configurational positioning play an important role in imparting RNA recognition specificity for IMP1 and IMP3.

**Figure 7.**
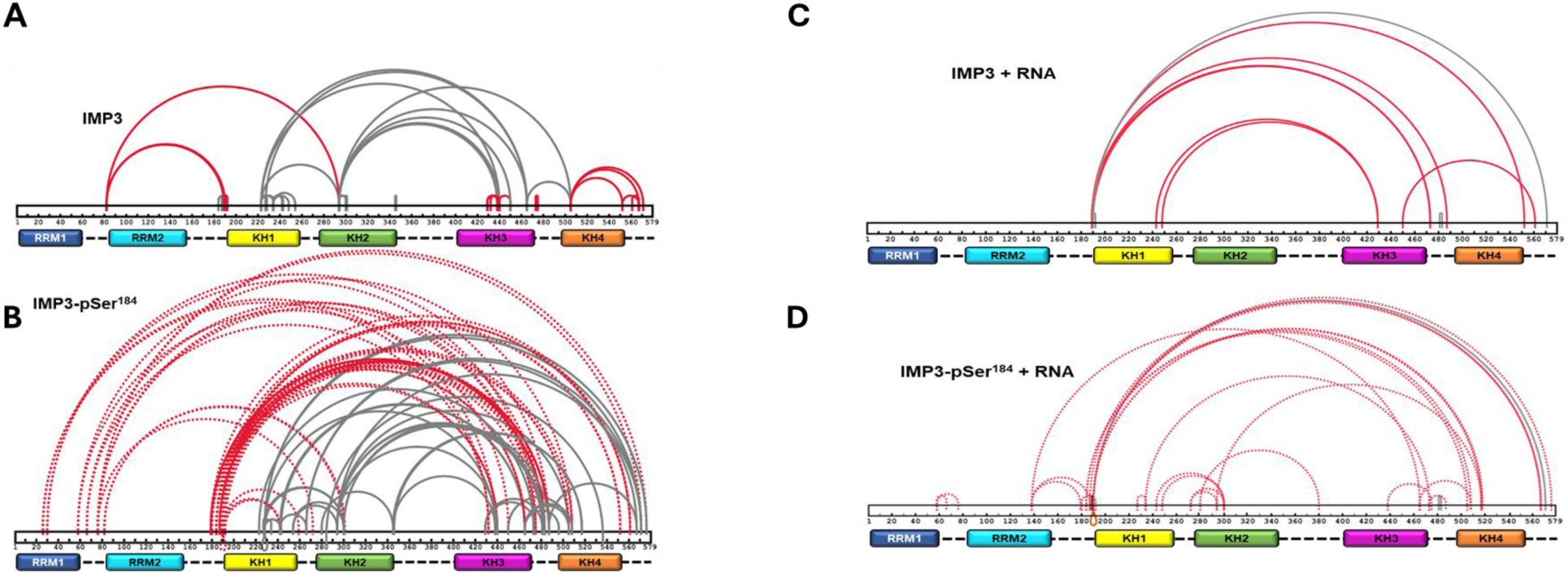
Cross-linking mass spectrometry analysis of IMP3 and IMP3-pSer^184^ reveal substantial configurational rearrangements upon phosphorylation. Crosslink maps of **A)** IMP3 and **B)** phosphorylated IMP3 (IMP3-pSer^184^) in the absence or **C-D)** presence of ANKRD RNA are shown. Cross-links identified using the amine-reactive cross-linker BS3 are displayed. Arcs denote cross-linked residues along the primary sequence, with the domain architecture of IMP3 (RRM1–2 and KH1–4) shown below each map. The crosslinking data collected in the absence and presence of RNA were analyzed as pairs. Red arcs indicate cross-links unique to the indicated condition, whereas gray arcs represent cross-links shared between the pair in each scenario. Phosphorylation at Ser184 results in a pronounced redistribution of inter-domain cross-links, consistent with large-scale configurational rearrangements that are further modulated by RNA binding.

### Computational modeling of structural changes induced by phosphorylation

To characterize the conformational ensembles of IMP1 and IMP3 in both their WT and phosphorylated states, consistent with the experimental crosslinking data, we performed coarse-grained molecular dynamics simulations using OpenMM. Proteins were represented at one-bead-per-residue resolution, with each bead centered on the Cα atom. Folded domains were stabilized in their native conformations by structure-based Gō-like interactions, whereas unstructured regions were modelled as flexible regions. Experimentally determined crosslinks were incorporated as finite-range, flat-bottom attractive interactions between the corresponding residues. For each protein, a 2.5 μs simulation in the absence of crosslink restraints was first performed to generate a diverse ensemble of high-entropy conformations. After discarding the first 20% of the trajectory, 200 conformations were selected as independent starting structures. Crosslink restraints corresponding to each experimental condition were then applied, followed by 60 ns of relaxation at 300 K and simulated annealing from 300 K to 0 K in 25 K increments, with 10 ns of simulation at each temperature. The resulting 200 structures constituted the conformational ensemble used for subsequent analyses. Full details of the coarse-grained model, interaction potentials and simulation protocol are provided in the Methods.

Coarse-grained simulations revealed distinct phosphorylation-dependent changes in the conformational ensembles of IMP1 and IMP3 (**Figure 8**). Phosphorylation increased the radius of gyration of IMP1, whereas IMP3 became more compact (**Figure 8A**). Analysis of the distances between the center of mass of the three structured dimers revealed that these changes arise from distinct rearrangements of their relative domain organization (**Figure 8B**). In IMP1, phosphorylation primarily led to an increased distance between the RRM1/2 and KH3/4 dimers, while the other interdimer distances remained comparatively unchanged. Phosphorylation of IMP3, in contrast, markedly decreased the distances between all three RRM1/2↔KH1/2, RRM1/2↔ KH3/4, and KH1/2↔KH3/4 dimers. Representative conformations illustrate these phosphorylation-dependent reorganization and the resulting expansion of IMP1 and compaction of IMP3 (**Figure 8C**).

**Figure 8.**
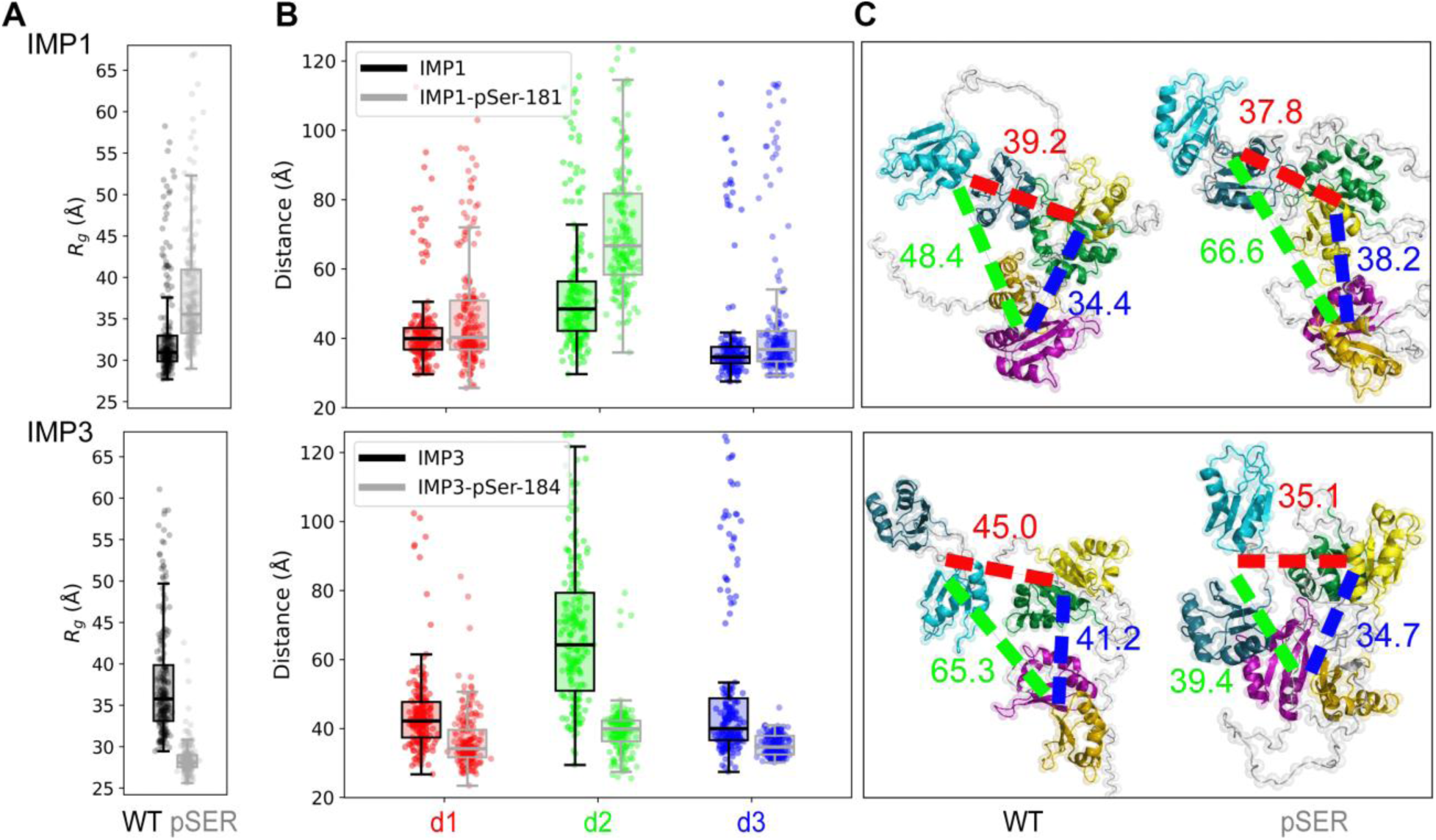
Phosphorylation differentially reorganizes IMP1 and IMP3 conformational ensembles. **A**) Radius of gyration (*R_g_*) of simulated WT and phosphorylated (pSER) IMP1 and IMP3 ensembles. **B**) Distributions of the three inter-dimer COM-COM distances (d1: RRM1/2 ↔ KH1/2; d2: RRM1/2 ↔ KH3/4; d3: KH1/2 ↔ KH3/4) reveal phosphorylation-dependent changes in domain organization. **C**) Representative WT and pSER conformations, with corresponding inter-dimer distances (Å) indicated by dashed lines. (d1: red; d2: green; d3: blue).

### Differences in end-to-end RNA distances reflect changes induced by phosphorylation

The XL-MS data reveal that the RNA binding domains and IDLs adopt different configurations upon phosphorylation and in the absence or presence of RNA. However, whether the structure of RNA is modulated by the differing configurations of the protein is not known. To address this question, we performed CW-EPR and double electron-electron resonance (DEER) spectroscopy^59,60^, the latter of which requires incorporation of two spin labels into the RNA oligonucleotide. Due to the challenge in synthesizing long RNA oligonucleotides, we performed this experiment only with IMP1, IMP1-pSer^181^ and its cognate 54-mer Zipcode RNA. 2′-aminouridine was introduced at two defined positions close to the 5′ and 3′ ends of the RNA during chemical synthesis of the oligonucleotides and post-synthetically spin-labeled with an isothiocyanate derivative of an isoindoline nitroxide (**Figure 9 & Supplementary Table 1**)^61^. The distance between the two spin centers was measured using DEER spectroscopy in the absence and presence of IMP1/IMP1-pSer^181^.

**Figure 9.**
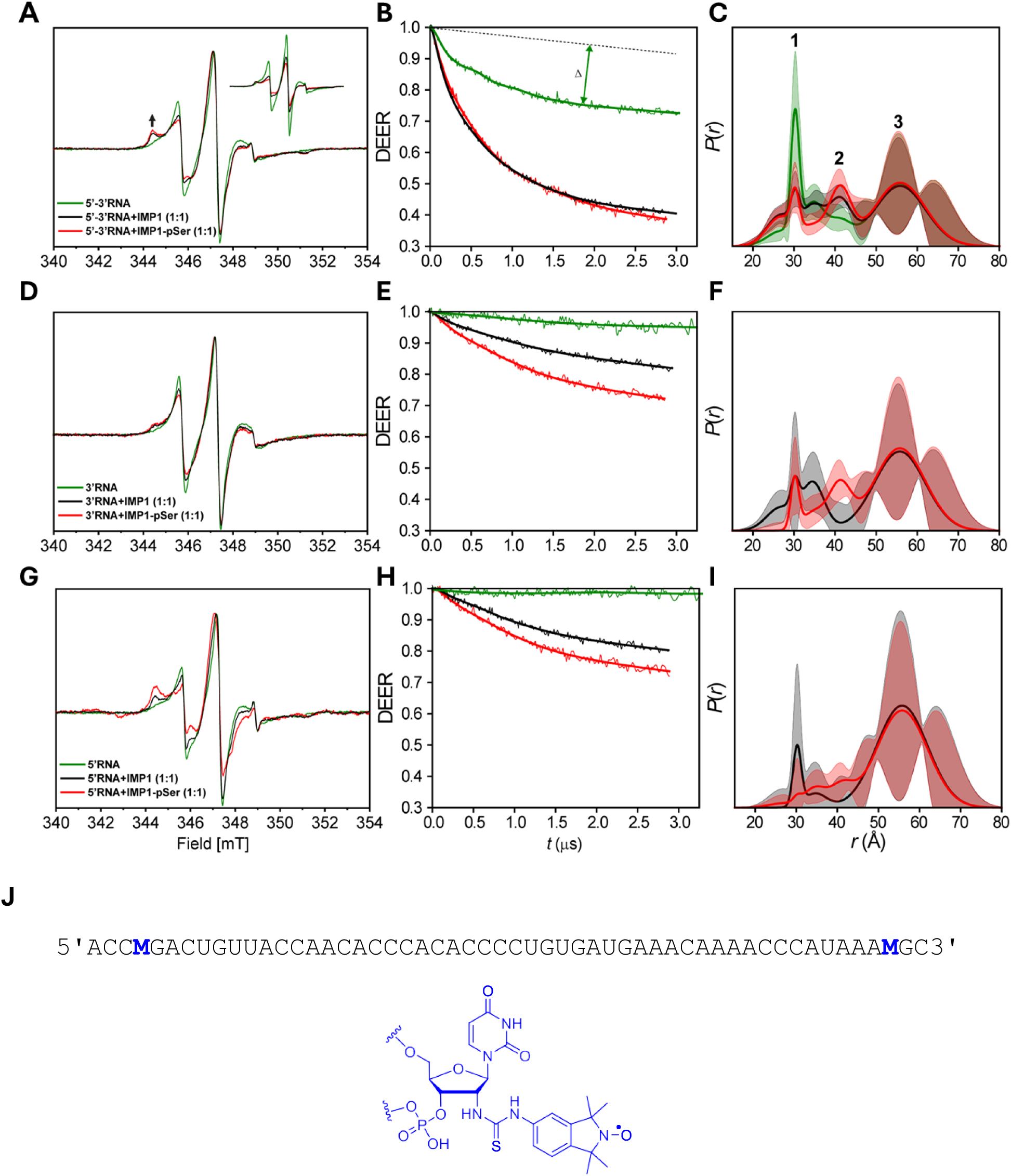
CW EPR and DEER spectroscopy of IMP1 and IMP1-pSer181 bound to Zipcode RNA. **A)** CW EPR spectra of doubly-labeled RNA in the absence (green) and presence of IMP1 (black), or and IMP1-pSer^181^ (red). **B)** Raw DEER decays and fits are presented for the experimentally determined distance distributions *P(r)*. **C)** Confidence bands (2σ) are shown about the best fit lines. CW EPR and DEER spectroscopy of singly labeled (**D-F**) 3ʹ-RNA and (**G-I**) 5′-RNA in the absence and presence of IMP1 or IMP1-pSer^181^. **J)** Zipcode RNA sequence and spin labeling positions. M denotes the position of the spin label.

Spin-label dynamics, inferred directly from the room-temperature CW EPR spectrum of the Zipcode RNA, report on molecular tumbling, local steric constraints, and backbone motions at the labeling sites. The observed intermediate-to-fast mobility of the spin labels in the free Zipcode RNA indicates that the RNA populates a partially folded and structurally ordered state (**Figure 9A-C**). Consistent with this interpretation, DEER measurements of the RNA alone resolve a folded subpopulation with a mean distance of ∼30 Å (population 1; **Figure 9C**). Binding to IMP1 alters the conformational ensemble of the Zipcode RNA (**Figure 9A-C**). The CW EPR spectral line shape shows increased immobilization upon IMP1 binding, consistent with reduced local mobility at the spin-labeled sites and slower overall molecular tumbling (**Figure 9A**). Interestingly, upon binding to IMP1, and even more prominently for IMP1-pSer^181^, a broader and longer-distance population (∼41 Å, population 2; **Figure 9C**) appears in the DEER distance distributions, indicating conformational heterogeneity within the bound RNA ensembles. The broadest population, centered near ∼56 Å (population 3; **Figure 9C**), is populated similarly under different conditions. IMP1 binding substantially increases the modulation depth (Δ) of the DEER traces beyond that of the doubly spin-labeled free RNA (**Figure 9B**). A higher modulation depth indicates a higher concentration of interacting spin pairs, which is essential for quantifying monomer versus dimer populations in a mixture. This suggests that multiple RNA molecules bind to IMP1. These data are consistent with mass photometry measurements that also reveal an ensemble of monomeric and dimeric IMP1 states (**Figure 5**).

To better dissect the intra- and inter-RNA distance contributions, we performed CW EPR and DEER spectroscopy on singly-labeled RNA (**Supplementary Table 1**). In these experiments, a DEER signal is observed only if two RNA molecules are bound to either a IMP1 monomer or oligomer. Interestingly, CW EPR revealed that the 3′ end of the Zipcode RNA is locally more flexible than the 5′ end (**Figures 9D & G**). In addition, the 5′ end becomes locally immobile when bound to IMP1-pSer^181^. DEER data on singly-labeled RNA also support the binding of two RNA molecules to IMP1, and this signal is enhanced for IMP1-pSer^181^. Considering the mass photometry data, it is highly likely that two RNA molecules bind to dimeric IMP1. Since IMP1 is predominantly a monomer in solution, these data suggest that IMP1-RNA interactions must occur via both *cis* and *trans* interactions where a select subset of the RBDs interact with one RNA molecule and the others engage the second RNA. Upon phosphorylation, the DEER data for the 3′ ends indicate more flexible relative orientation sampling populations 2 and 3 (**Figures 9E-F**), whereas the 5′ ends are more distant (i.e., population 3) and less flexible relative to each other (**Figures 9H-I**). Thus, populations 2 and 3 mainly represent intermolecular distance contributions from the 3′ and 5′ ends, respectively. Additionally, a decrease in population 1 indicates that IMP1 binding partially promotes unfolding and extension of the RNA. Generally, IMP1-pSer^181^ imposes greater restriction on monomeric/dimeric RNA dynamics than the IMP1 protein (**Figures 9D-I**). Overall, both the CW EPR and DEER data suggest that the domains interact differentially with RNA upon phosphorylation, and that the structure of the RNA is different when bound to the phosphorylated versus unmodified form of IMP1. These data are in strong agreement with both the XL-MS and NMR results.

In addition, we tested the effects of IMP3 or IMP3-pSer^184^ interactions with the Zipcode RNA and observed overall CW-EPR behavior similar to that of IMP1 and IMP1-pSer^181^ (**Supplementary Figure 3A**). These data suggest that there is not an overall difference in RNA binding affinities of both proteins to RNA. However, DEER measurements indicated that the phosphorylation-dependent changes observed for IMP1 were attenuated in IMP3, with RNA-bound IMP3-pSer^184^ exhibiting broader distance distributions and a reduced population 2 (**Supplementary Figures 3B & C**). These data suggest that the conformations of the bound RNA are different between IMP1 and IMP3, and likely induced by the configurational changes in the two proteins induced upon their respective pSer modifications.

## Discussion

Post-translational modification of intrinsically disordered regions (IDRs) has emerged as a pervasive mechanism for regulating RNA-binding proteins (RBPs), yet how such modifications reshape RNA recognition and functional specificity at a structural level remains largely unresolved. Here, we show that phosphorylation by mTORC2 within a disordered linker of IGF2BP paralogs operates as a configurational switch, reorganizing long-range domain arrangements without perturbing folded RNA-binding domains or strongly altering RNA-binding affinity. Despite sharing >74% sequence identity, IMP1 and IMP3 display distinct phosphorylation-dependent and independent configurational landscapes, providing a structural basis for their paralog-specific RNA regulatory functions.

Phosphorylation at Ser181 in IMP1 and Ser184 in IMP3 causes only moderate reductions in RNA-binding affinity, consistent with the multivalent architecture of these proteins (**Figure 1**). Each IGF2BP contains six RNA-binding domains (RBDs) whose collective engagement can tolerate perturbations at individual contacts without loss in overall RNA binding. This finding agrees with previous studies showing that phosphomimetic mutations at Ser181 or Src kinase-targeted Tyr396 do not significantly alter steady-state RNA binding across diverse transcripts, despite pronounced functional consequences^45^. Together, these observations indicate that regulation of IGF2BPs cannot be inferred from affinity measurements alone. Instead, our data demonstrate that phosphorylation leaves the secondary structure, thermal stability, and folding of RRM and KH domains intact, while redistributing interdomain configurations through the intrinsically disordered linker connecting RRM2 and KH1. NMR, CD, and nanoDSF analyses converge on a model in which phosphorylation locally perturbs the disordered region and alters its interactions with folded domains, without inducing global conformational changes. Our findings establish the linker as an active regulatory element that transduces post-translational signals, in this case phosphorylation, into large-scale architectural rearrangements.

Our data also highlight that homologous phosphorylation events in IMP1 versus IMP3 elicit distinct configurational responses. In IMP1, phosphorylation loosens interdomain packing and increases configurational heterogeneity, correlating with enhanced accessibility for oligomerization. In contrast, phosphorylation of IMP3 drives a more compact architecture characterized by increased intramolecular crosslinks. These changes are captured and revealed through a combination of crosslinking mass spectrometry and MD simulation analysis (**Figures 6-8**). These divergent outcomes occur despite near-identical domain organization and underscore how subtle differences in IDR sequence can encode paralog-specific regulatory behavior. Thus, phosphorylation does not impose a uniform structural outcome but instead reshapes energy landscapes intrinsic to each protein. RNA binding further amplifies these differences. Both paralogs dimerize upon RNA engagement, consistent with cooperative assembly on clustered RNA elements. However, phosphorylation alters the stability and homogeneity of the resulting complexes, particularly for IMP3, where phosphorylated protein forms less uniform RNA-bound assemblies. These observations suggest that RNA serves not only as a ligand but also as a conformational selector, stabilizing particular protein architectures depending on the modification state. Importantly, phosphorylation-dependent configurational changes propagate to the RNA itself. DEER and CW-EPR spectroscopy reveal that IMP1 binding remodels RNA end-to-end distance distributions and that phosphorylation biases the RNA ensemble toward more extended and constrained configurations without eliminating structural heterogeneity. Rather than enforcing a single RNA fold, phosphorylated IMP1 shifts the population of pre-existing states. Such ensemble-level modulation may influence the accessibility of regulatory elements, accommodate additional binding partners, or alter the geometry of higher-order ribonucleoprotein assemblies. Interestingly, the changes in RNA conformations are also unique to IMP1 and IMP3 and their pSer variants. These differences likely explain the origins of functional diversity and outcomes that occur in the cell when the same RNA is bound by these structural paralogs.

From a mechanistic standpoint, IGF2BP paralogs do not possess intrinsic catalytic activity; instead, they function as high-affinity, multivalent RNA binders that scaffold messenger ribonucleoprotein (mRNP) assemblies. This non-enzymatic mode of action implies that functional specificity cannot arise from chemical transformation of RNA but must instead be encoded through spatial and temporal organization and access of RNA-protein interfaces. In this framework, phosphorylation-induced configurational rearrangements provide a compelling mechanism for regulation. By reorganizing interdomain geometry without substantially altering global RNA-binding affinity, IGF2BP proteins can modulate which regions of a bound mRNA are exposed or occluded to other RNA-regulatory factors, including translation initiation components, decay machineries, and remodeling enzymes. Such configurational gating offers a means to differentially regulate identical RNA targets across cellular contexts. The distinct phosphorylation-dependent architectures observed for IMP1 and IMP3 therefore suggest that paralog specificity arises not just from differential RNA recognition per se, but from how each protein positions RNA within heterogeneous mRNP environments. Consistent with this view, IGF2BP1 and IGF2BP3 have been implicated in divergent regulatory outcomes on shared transcripts, including opposing effects on translation and stability, despite binding overlapping RNA elements^51^.

Configurational control of nucleic acid binding specificity via interdomain rearrangements is an established regulatory principle in multidomain nucleic acid binding proteins. A well-characterized example is Replication Protein A (RPA), a multidomain single-stranded DNA-binding protein in which interdomain configurations dictate nucleic acid binding and downstream activities. We have shown that such configurations are dynamically tuned by single- and multisite phosphorylation, which modulates both DNA engagement^62^ and protein-protein interactions, and can reciprocally regulate kinase activity itself through a feedback mechanism^63,64^. In RNA-binding proteins, contextual features such as single-versus double-stranded RNA are predicted to further bias domain arrangements and thereby influence regulatory outcomes^65^. Analogous to RPA, IGF2BP paralogs lack intrinsic catalytic function and instead encode regulatory specificity through phosphorylation-sensitive reorganization of domain geometry. The configurational tuning of IMP1 and IMP3 described here therefore represents one component of a broader arsenal of mechanisms, in which post-translational modification reshapes conformational ensembles to bias molecular access to RNA or binding sites on the protein. These sophisticated structural changes in turn are likely to contribute to paralog-specific RNA regulation.

Together, our findings support a model in which IGF2BP paralogs function as architectural regulators of RNA. mTORC2-dependent phosphorylation acts as a configurational switch that tunes interdomain arrangements through intrinsically disordered regions, thereby modulating how bound mRNAs are presented to downstream effectors involved in translation, stability, remodeling, and condensate assembly. In this view, functional regulation emerges not from changes in intrinsic RNA-binding strength, but from phosphorylation-dependent redistribution of conformational ensembles that govern molecular access and assembly within mRNPs. More broadly, these findings add to the growing recognition that phosphorylation of IDPs acts as a global regulatory mechanism, reshaping conformational ensembles and interdomain communication to encode specificity, plasticity, and context-dependent functional outputs in proteins and protein-nucleic acid complexes^66,67^.

## Acknowledgements

This work was supported by funding from the National Institutes of Health - National Institute of General Medical Science R35-GM149320 (E.A.), R01-GM143179 (S.O.), R35-GM158220 (H.A.); the National Cancer Institute R01-CA305375 (E.A.), R01-CA265877 (R.D.); the Office of the Director S10-OD030343 (E.A.), S10-OD036230 (R.D.); the Icelandic Research Fund (grant no. 206708; S.Th.S) and a PhD fellowship from the University of Iceland Research Fund (P.R.). Mass Spectrometry analyses for pSer identification were performed by the Mass Spectrometry Technology Access Center at McDonnell Genome Institute (MTAC@MGI) at Washington University School of Medicine. This work was aided by the GCE4All Biomedical Technology Development and Dissemination Center supported by National Institute of General Medical Science grant RM1-GM144227. We thank Maxim Drömer for testing the feasibility of isotopic labeling of IMP1. We thank Ethan J. Hasenoehrl for assistance with XL-MS experiments.

DOI for the code repository for MD simulations: https://doi.org/10.5281/zenodo.21923103

## Author Contributions

Conceptualization: EA and SO

Methodology: VK, VS, JM, MTL, RT, IE, RC, RK, AV, PR, RAM, RBC, STS, YL,RD, HA, BBB, SO & EA

Investigation: VK, VS, JM, MTL, RT, IE, RC, RK, AV, & PR

Visualization: VK, VS, RT, HA, MTL, RK, BBB, YL, RD and EA

Funding acquisition: EA, SO, HA, RD, YL, STS, & RAM

Project administration: EA

Supervision: STS, YL, RD, HA, BBB, SO, and EA

Writing – original draft: EA, RD, RT, YL & HA

Writing – review & editing: all authors

## Methods

### Plasmids and RNA oligonucleotides

Plasmids for protein overexpression were generated by Genscript Inc. Briefly, codon-optimized open reading frames (ORFs) for IMP1 or IMP3 were synthesized with a N-terminal poly-his tag, a SUMO-tag, and followed by a SUMO protease cleavage site. An additional C-terminal poly-His tag was engineered to separate truncated proteins during non-canonical amino acid incorporation. These ORFs were subcloned into pRSF-Duet plasmids. TAG codons for site-specific incorporation of Phosphoserine (pSer) were introduced using site-directed mutagenesis.

Unlabeled RNA oligonucleotides were purchased from Integrated DNA Technologies. The substrate used for IMP1 (oligo-1) was a 54-nucleotide (nt) *cis*-acting element, termed Zipcode, that is positioned directly following the termination codon in the 3′ untranslated region (UTR) of β-actin mRNA^42,54^. The substrate used for IMP3 (oligo-2) is a 121-nt region in exon 29 from ANKRD17 mRNA^33,55^. For the synthesis of the doubly spin-labeled 54-nucleotide Zipcode RNA, 2′-aminouridine nucleotides were incorporated at specific sites and post-synthetically spin labeled with an isothiocyanate derivative of an isoindoline nitroxide as described^61^. Sequences for the two substrates are detailed in **Supplementary Table 1**.

### Expression and purification of wild type IMP1 and IMP3

Human IMP1 and IMP3 wild type proteins were recombinantly produced using plasmid pRSFDuet1-IMP1 and pRSFDuet1-IMP3, respectively, and expressed in BL21(DE3) *E. coli* cells. Transformants were selected using kanamycin (50 µg/mL). An overnight starter culture was prepared by inoculating single colony in 50 mL of Luria Broth (LB) media with kanamycin selection. 1% of the inoculum was added to 1 L of LB media and grown at 37 °C with shaking at 250 rpm, until the OD_600_ reached 0.6, followed by induction with 0.4 mM IPTG. Induction was carried out at 20 °C for 16-20 hrs. Cells were harvested and resuspended in 30 mL of IMP resuspension buffer (50 mM Tris-HCl, pH 8.0, 1 M NaCl, 10% glycerol, 1X protease inhibitor cocktail, and 1 mM tris(2-carboxyethyl)phosphine (TCEP)). Cell lysis was performed using 0.4 mg/mL lysozyme, stirred for 30 min at 4 °C, followed by sonication. The clarified lysate was then fractionated over a Ni^2+^-NTA agarose column (Gold Biotechnology Inc.) with a 2 M NaCl wash step to remove non-specifically bound nucleic acid contaminants. Protein was eluted using cell resuspension buffer containing 350 mM imidazole. Fractions containing pure protein were then pooled and digested with ULP1 protease (1:100) for 2 hrs at 4 °C to clip off the N-terminal His- and SUMO-tags. Digested protein was then diluted with H_0_ buffer (50 mM Tris-HCl, pH 8.0, 10% glycerol, 1X PIC, and 1 mM TCEP) to match the conductivity with H_100_ buffer (H_0_ + 100 mM NaCl) and further fractionated over a fast flow heparin column (Cytiva Inc.). Bound proteins were eluted using H_100_ buffer and a linear gradient (0.1 to 1M NaCl). Fractions containing protein were pooled and concentrated using an Amicon spin concentrator (10 kDa molecular weight cut-off). The concentrated protein was then fractionated over a S200 column (Cytiva Inc.) using Storage buffer (50 mM Tris-HCl, pH 8.0, 10% glycerol, 300 mM NaCl, and 1 mM TCEP). Purified protein was flash frozen using liquid nitrogen and stored at −80 °C. IMP1 and IMP3 concentrations were measured spectroscopically using ℇ_280_ = 28,880 and 25,900 M^-1^cm^-1^, respectively.

### Generation of site-specific phosphoserine carrying IMP1 (IMP1-pSer^181^) and IMP3 (IMP3-pSer^184^)

Phosphoserine (pSer) was site specifically incorporated at position S181 in IMP1 and S184 in IMP3, through genetic code expansion (GCE)^52,53^. Briefly, TAG stop codons were introduced at these positions in the ORF using site-directed mutagenesis. pRSFDuet1-IMP1-S181-TAG or pRSFDuet1-IMP3-S184-TAG plasmids were co-transformed with pKW2-EFSep (machinery plasmid for phosphoserine incorporation; Addgene Plasmid #173897) in BL21 (DE3) Δ*serB E. coli* competent cells and successful transformants were selected using chloramphenicol (25 μg/mL) and kanamycin (50 μg/mL) on Luria agar plates. About a dozen colonies were scraped and inoculated in ZY-non inducing media (ZY-NIM; **Supplementary Table 2**) to prepare a starter culture and grown overnight at 37 °C with shaking at 250 rpm. For the expression culture, 1% of the inoculum was added to ZY-auto induction media (ZY-AIM; **Supplementary Table 2**) and grown until the OD_600_ reached 1.5. The cultures were shifted to 20 °C and grown for an additional 20 hours. Cells were then harvested and resuspended in IMP cell resuspension buffer containing phosphatase inhibitors (50 mM Tris-HCl, pH 8.0, 1 M NaCl, 10% glycerol, 1X protease inhibitor cocktail, 1 mM TCEP, 50 mM sodium fluoride, 10 mM sodium pyrophosphate, and 1 mM sodium orthovandate). pSer carrying IMP1 and IMP3 proteins were purified similar to the wildtype protein. IMP1-pSer^181^ and IMP3-pSer^184^ protein concentrations were measured spectroscopically using ℇ_280_ = 28,880 and 25,900 M^-1^cm^-1^, respectively. Yields of pSer-containing IMP1/IMP3 were ∼4-5 mg/L of culture.

To obtain ^15^N isotope-incorporation in IMP1 samples for NMR experiments, protein was expressed in minimal medium enriched with 1 g/L ^15^NH_4_Cl and ^15^N-labeled Celtone^®^ (Cambridge Isotopes Laboratories Inc.). Site-selective incorporation of pSer at position S181 in conjunction with ^15^N-labeling was achieved as described^68^. Briefly, a high density culture (OD_600_ =3.7) of cells co-transformed with plasmids pRSFDuet1-IMP1-S181TAG and pKW2-EFSep in minimal medium enriched with 1 g/L ^15^NH_4_Cl and 2 g/L ^15^N-labeleld Celtone^®^ was prepared and protein expression was induced with 0.4 mM IPTG for 48 h. Proteins were purified as described above and buffer exchanged into 50 mM sodium phosphate buffer pH 6.5, 100 mM NaCl, 1 mM DTT, 0.5 mM EDTA spiked with 5% D_2_O before transfer into a 5 mm Shigemi tube.

### Analysis of pSer incorporation using Phos-Tag SDS-PAGE

Incorporation of pSer into IMP1 or IMP3 was assessed using Phos-tag SDS PAGE (Fujifilm Inc.). Phos-Tag SDS-PAGE gel (10%) was prepared by adding 0.05 mM of the Phos-tag reagent and 0.1 mM MnCl_2_ to the resolving gel (before pouring the gel). Other steps for preparation of the resolving and stacking gels, and conditions for running the gel, are as defined for standard SDS-PAGE analysis. Gels were stained using Coomassie stain. The protein carrying pSer runs at a higher apparent molecular weight in Phos-tag SDS PAGE analysis.

### Proteomics and data analysis

For detection of pSer incorporation by mass spectrometry, proteins were resolved by SDS-PAGE on a 10% polyacrylamide gel. Protein bands of interest were excised and subjected to in-gel digestion. Gel pieces were washed with 100 mM ammonium bicarbonate in acetonitrile (AmBic/ACN), reduced with 10 mM dithiothreitol at 50 °C for 30 min, and alkylated with 100 mM iodoacetamide for 30 min at room temperature in the dark. Following alkylation, gel pieces were washed with AmBic/ACN and incubated overnight at 37 °C with 2 µg trypsin. Peptides released during digestion were collected, and residual peptides were sequentially extracted from the gel pieces by incubation at room temperature with gentle shaking for 10 min in 50% ACN/5% formic acid, followed by 80% ACN/5% formic acid, and 100% ACN. All supernatants were pooled and lyophilized to completion using a SpeedVac concentrator. Peptides were reconstituted in 0.1% formic acid in water and analyzed by LC-MS/MS without phosphopeptide enrichment. Peptides were loaded onto a trap cartridge and separated on a PepMap RSLC C18 analytical column (75 µm × 15 cm, 3 µm particle size) using a 60-min linear gradient of solvent A (0.1% formic acid in water) and solvent B (0.1% formic acid in acetonitrile) on a Vanquish Neo UHPLC system coupled to an Orbitrap Eclipse Tribrid mass spectrometer (Thermo Fisher Scientific). Raw data were searched using Mascot (v2.8.0) against a custom database consisting of *Escherichia coli* (strain BL21(DE3)) proteins supplemented with two custom protein sequences. Phosphorylation of serine residues was included as a variable modification. Peptide and protein identifications were filtered to a 1% false discovery rate and reported using Scaffold (v5.1.1).

### Fluorescence anisotropy analysis of RNA binding

The RNA binding activity of both unmodified and phosphorylated IMP proteins was measured using fluorescence anisotropy. 25 nM of 5′-FAM-labeled Zipcode RNA (54 nt) or 5′-FAM-labeled ANKRD RNA (121 nt) were used to quantify binding to IMP1/IMP1-pSer^181^ or IMP3/IMP3-pSer^184^, respectively. RNAs were resuspended in RNA binding buffer (20 mM Tris-HCl pH 7.5, 100 mM NaCl, 1 mM MgCl_2,_ 1 mM TCEP-HCl, and 5% glycerol). 1.2 mL of this working RNA stock was added to a 10 mm path length quartz cuvette (Starna Cells Inc.) while maintaining the temperature at 23°C. Fluorescence anisotropy of 25 nM 5′ FAM-labeled RNA with increasing concentrations of protein was measured using a PC1 spectrofluorometer (ISS Inc.) and data collection was performed using Vinci3 software associated with the instrument. In general, samples were excited at 488 nm and emission was recorded using 520 nm band pass emission filter. Five consecutive anisotropy readings were acquired for RNA alone and with each different concentration of protein added. The concentrations of RNA, proteins, and reduction in fluorescence intensity were corrected for effects due to dilution resulting from the stepwise addition of the protein. Proteins binding to fluorescein-labeled molecules usually results in fluorescence quenching, because the fluorescence quantum yield of the bound species is lower than that of free RNA. If it is not corrected, this artifact can result in significant errors in measuring binding affinity. Further, measured anisotropy and any change in fluorescence intensity values were also corrected. Quantum-yield corrections were made using Equation 1 where: *A_c_*= corrected anisotropy, *A*= measured anisotropy, *A_f_*= anisotropy of the free 5′-FAM-labeled Zipcode or ANKRD RNA, *A_b_*= anisotropy of the bound 5′-FAM-labeled Zipcode or ANKRD RNA, *Q_f_*= fluorescence quantum yield of the free 5′-FAM-labeled Zipcode or ANKRD RNA, and *Q_b_*= fluorescence quantum yield of the bound 5′-FAM-labeled Zipcode or ANKRD RNA.

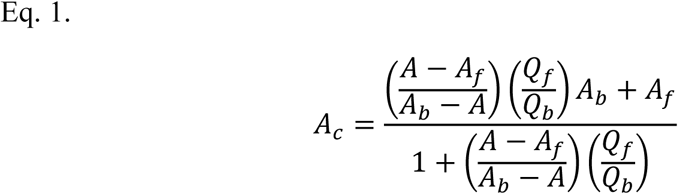

The anisotropy data were fit to a mass-action binding isotherm with ligand depletion using Equation 2 where *Y*=measured anisotropy, *r*_min_= anisotropy of free 5′-FAM-labeled Zipcode or ANKRD RNA, *r*_max_= anisotropy of fully bound 5′-FAM-labeled Zipcode or ANKRD RNA, *E*= total IMP1 (p) or IMP3 (p) concentration, *X*= total 5′-FAM-labeled Zipcode or ANKRD RNA concentration, and *K_D_*= dissociation constant.

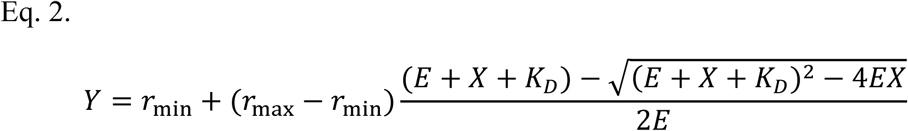

Anisotropy values (Y) versus protein concentration (X) data points were fit keeping *r*_min_, *r*_max_, and *K_D_* as free parameters using non-linear least squares regression. Finally mean ± SEM was estimated, and data was plotted using Graph pad prism 9.

### Secondary structure analysis using circular dichroism

CD analysis for wild type and phosphorylated proteins was performed using a Chirascan V100 spectrophotometer (Applied Photophysics Inc.). Proteins were buffer-exchanged with the CD buffer (5 mM Tris-HCl pH 7.5, 100 mM NaF, and 1 mM TCEP-HCl) before the measurements. A nitrogen-fused set up with a Type-0590 cuvette (Starna Scientific Inc; cell path of 0.2 mm) was used to perform the experiments at 20 °C. All CD traces were recorded between 200-260 nm, and traces were background-corrected using the CD buffer. 3 µM of IMP1, IMP1-pSer^181^, IMP3, and IMP3-pSer^184^ were used, and 10 scans were collected for each sample and averaged. CD data were analyzed using BestSel^69^ to obtain secondary structure calculations. Mean ± SEM was estimated from three independent measurements. Data after normalization was plotted in Graph Pad prism 9.

### Mass photometry analysis of oligomeric states

All measurements were carried out on a TwoMP mass photometry instrument (Refeyn Ltd.) as previously described^62^. Glass coverslips (No. 1.5H thickness, 24 × 50 mm; VWR) were cleaned by sonication in isopropanol, rinsed thoroughly with deionized water, and dried under a stream of nitrogen gas. For each measurement, a clean coverslip was mounted on an oil-immersion objective lens (Olympus PlanApo N, 60X, 1.42 NA), and a holey silicone gasket (Refeyn Ltd.) was adhered to the upper surface of the coverslip. All dilutions and measurements were performed at room temperature (23 ± 2 °C) in 1X Mg^2+^/Ca^2+^ buffer (20 mM HEPES, pH 7.5, 150 mM KCl, 5 mM MgCl_2_, 5 mM CaCl_2_) supplemented with 1 mM DTT. Samples of IMP1, IMP3, and their phosphoserine variants, as well as protein-RNA mixtures, were equilibrated at 23 ± 2 °C for 5 min, after which 1 µL of each sample was rapidly diluted into 15 µL of buffer and immediately measured. Particle landing events were video-recorded for 1 min. Individual interferometric scattering events corresponding to single particles landing on the coverslip surface were identified and analyzed. A protein mass standard (β-amylase; Sigma A8781-1VL) was used to generate a contrast-to-mass calibration curve. Event-based histograms of calibrated molecular mass were compiled from all detected landing events acquired during the 1-min recording. Histogram peaks were fit using nonlinear least-squares Gaussian fitting to determine the mean molecular mass and associated uncertainty. Reported uncertainties correspond to the standard deviation obtained from Gaussian fitting.

### NMR spectroscopy

^15^N-TROSY-HSQC spectra of IMP1 (100 µM), IMP1-pSer^181^ (165 µM), IMP1 with Zipcode RNA (50 µM) at ratio 1:1 and IMP1-pSer^181^ with Zipcode RNA (100 µM) at ratio 1:1 were recorded on a Bruker 700 MHz spectrometer equipped with a cryogenically cooled TCI probe head. In the indirect ^15^N-dimension, the carrier was placed at 117 ppm, spectral width was set to 33 ppm and signal was acquired for 51 ms. The number of scans was set to 300 (IMP1) and 260 (IMP1-pSer^181^) with the recycling delay set to 1 sec. The experiments was recorded at 298K. NMR data were processed using NMRPipe^70^ and analyzed and visualized using CCPNMR Analysis^71^. The contour level in the overlay plots are adjusted to account for differences in concentration and signal averaging.

### Cross-linking mass spectrometry (XL-MS) analysis

IMP1, IMP3, IMP1-pSer^181^, or IMP3-pSer^184^ (10 µM) were incubated in the absence or presence of cognate RNA substrates (10 µM) with 5 mM bis(sulfosuccinimidyl) suberate (BS3) at room temperature for 15 min in a final volume of 20 µL reaction buffer (50 mM HEPES, pH 7.8, 100 mM KCl, 10% glycerol). Cross-linking reactions were quenched by addition of 2 µL of 1 M ammonium acetate and incubated for 15 min at room temperature. Samples were subsequently resolved by SDS-PAGE. Protein bands were excised and destained using a solution of 50 mM ammonium bicarbonate and 50% acetonitrile. Gel pieces were reduced with 100 mM dithiothreitol in 25 mM ammonium bicarbonate for 30 min at 56 °C, followed by alkylation with 55 mM iodoacetamide in 25 mM ammonium bicarbonate for 25 min at room temperature in the dark. Gel pieces were washed with 50 mM ammonium bicarbonate and 50% acetonitrile, dehydrated with 100% acetonitrile, and rehydrated with sequencing-grade trypsin (0.6 µg; Promega). Digestion was carried out overnight at 37 °C. Peptides were extracted using 10 µL of 50% acetonitrile and 0.1% formic acid, transferred to fresh microcentrifuge tubes, vortexed for 5 min, and centrifuged at 15,000 rpm for 30 min. Extracted peptides were dried, reconstituted in 0.1% formic acid in water, and injected onto a Neo trap cartridge coupled to an analytical column (75 µm inner diameter × 50 cm, PepMap Neo C18, 2 µm particle size). Peptide separation was performed using a 120-min linear gradient of solvent A (0.1% formic acid in water) and solvent B (0.1% formic acid in acetonitrile) on a Vanquish Neo UHPLC system coupled to an Orbitrap Eclipse Tribrid mass spectrometer equipped with a FAIMS Pro Duo interface (Thermo Fisher Scientific). MS/MS data were searched for cross-linked peptides using Proteome Discoverer v3.0 with the XlinkX node against a database consisting of the target protein sequence(s). BS3 specificity was defined for lysine residues and protein N-termini. Cross-link identifications were filtered to a 1% false discovery rate. Cross-linking data were visualized using xiVIEW^72^ and figures were prepared using Inkscape.

### Nano-DSF and thermal stability analysis

Thermal unfolding of IMP1, IMP3 or IMP1-pSer^181^, or IMP3-pSer^184^ were performed with a Prometheus NT.48 instrument (NanoTemper Technologies), which measures the intensity of the intrinsic protein fluorescence at 330 and 350 nm across a temperature gradient. All proteins were prepared at a concentration of 4 µM in buffer (50 mM Tris-Cl, pH 8.0, 300 mM NaCl, 1 mM TCEP-HCl) and loaded into standard capillaries (NanoTemper Technologies) for analysis. All experiments were conducted with a linear thermal ramp from 20 to 95 °C using a heating rate of 2 °C/min. The fluorescence intensity ratio (F350/F330) was plotted against temperature, and the inflection point (IP350/330) of the transition was derived from the maximum of the first derivative for each measurement using PR ThermControl Software v. 2.1.5 (NanoTemper Technologies). All experiments were carried out in triplicate. Mean and standard deviation were calculated from three independent measurements.

### Continuous wave (CW)-EPR and DEER spectroscopy

CW-EPR spectra of spin-labeled Zipcode RNA (**Supplementary Table 1**) samples in the absence or presence of IMP1 or IMP1-pSer^181^ were collected at room temperature on a Bruker EMX spectrometer operating at X-band frequency (9.5 GHz) using 2 mW incident power and a modulation amplitude of 1 G. DEER spectroscopy was performed on a Fathom EPR Spectrometer (High Q Technologies Inc.) operating at X-band frequency (9.6 GHz) with the DEER-Stitch method combining pulse sequences (16 steps phase cycling) at 50 K. Pulse lengths were 15 ns (π/2) and 30 ns (π) for the Gaussian probe pulses and 200 ns for the Adiabatic pump pulse. The frequency separation was 80 MHz. Samples for DEER analysis were cryoprotected with 30% (vol/vol) glycerol and flash-frozen in liquid nitrogen utilizing a SQBA™ sample cartridge. Primary DEER decays were analyzed using a home-written software (DeerA, Dr. Richard Stein, Vanderbilt University) operating in a Matlab (MathWorks) environment as previously described^73^. Briefly, the software carries out analysis of the DEER decays obtained under different conditions. The distance distribution is assumed to consist of a sum of Gaussians, the number and population of which are determined based on a statistical criterion.

### Computational structural modelling of IMP protein system

To study the conformational landscape of the IMP protein system following experimental crosslinking (XL-MS), we performed coarse-grained molecular dynamics simulations. All simulations were performed using the OpenMM molecular dynamics engine, with a Langevin middle integrator. Each protein was represented using a one-bead-per-residue coarse-grained model in which consecutive residues within the protein were connected by harmonic bond and angle potentials. The equilibrium bond lengths (*r*_0_) and bond angles (*θ*_0_) were obtained directly from the input reference structure. Harmonic bond and angle potentials were defined as *E_b_* = *k_b_*(*r* − *r*_0_)^2^ and *E_θ_* = *k_θ_*(*θ* − *θ*_0_)^2^ where (*r*) and (*θ*) are the instantaneous bond length and bond angle, respectively. Bond force constants were set to *k_b_* = 20 · 10^3^ *kJ mol*^−1^*nm*^−1^. To account for the different flexibilities of structured and unstructured regions, harmonic angles located within folded domains were assigned a force constant of *k_θ_* = 40 *kJ mol*^−1^*rad*^−1^, whereas angles centered on unstructured regions were softened *k_θ_* = 10 *kJ mol*^−1^*rad*^−1^, allowing greater flexibility. Dihedrals angles were added only within structured regions, using a periodic torsion potential assigned only when the central bond of a four-residue segment was entirely contained within a structured domain. The equilibrium dihedral angles were extracted from the reference structure, and each retained torsion was represented using an energy term of the form, *E_ϕ_* = *k_ϕ_*(1 + cos(*nϕ*− *ϕ*_0_) where *ϕ* is the instantaneous dihedral angle and *ϕ*_0_ is the reference value. The first harmonic was assigned a force constant of 1.0, while the third harmonic was assigned half this strength. The boundaries of the structured and unstructured domains were obtained from the UniProt database (IMP1: Q9NZI8; IMP3: O00425).

The native structure of the folded domains was maintained using Gō-like attractive interactions between residue pairs identified from the reference atomistic structure with the Shadow contact algorithm. These native contacts were modeled with 12-10 Lennard-Jones potentials of the form 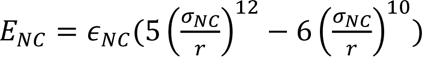, where (*σ*_NC_) is the native inter-residue distance extracted from the reference structure, and (*ε_NC_*) is the well depth set to 12.5 *kJ mol*^−1^. No native contacts were assigned to unstructured regions. The KH-KH2 and KH3-KH4 dimers were maintained by introducing native contacts between residues belonging to the two monomers, following the same procedure used for intradomain native contacts. No native contacts were assigned between the RRM1 and RRM2 domains. Instead, their separation was maintained using a flat-bottom center of mass restraint, allowing the domains to translate and rotate freely while preventing their center-of-mass distance from exceeding the initial value in the reference structure. Excluded volume interactions were added between all non-excluded residue pairs, through a purely repulsive Weeks-Chandler-Andersen (WCA) potential, where the specific residue-pair excluded volume diameter (*σ_ij_*) was obtained from the HPS force field and the repulsive energy scale was set to 0.2 *kJ mol*^−1^(**Supplementary Figure 4**).

Experimentally derived crosslinks were represented as finite-range attractive flat-bottom potential between the corresponding residue beads. For each crosslinked pair, the interaction energy was defined as

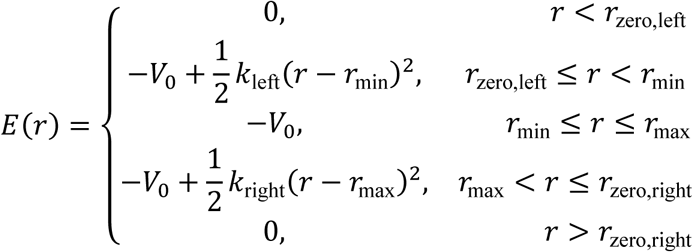

where *V*_0_ = −7.5 *kJ mol*^−1^ is the well depth, and was chosen to allow reversible binding at 300K. We define the flat attractive region between *r_min_* = 1.5 *nm* and *r_max_* = 2.5 *nm*. *r*_zero,left_ and *r*_zero,right_ define the distances at which the restraint smoothly returns to zero and were set to 1.4 *nm* and 3.0 *nm*, meaning distance between 2.5 and 3.0 nm are possible but energetically penalized. The harmonic force constants were calculated for left and right side by:

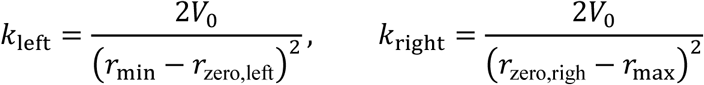

To generate representative conformational ensembles consistent with each experimental crosslink dataset, simulated annealing was performed from a diverse set of high-entropy initial conformations. First, a 2.5 μs simulation of the protein without crosslinks was carried out using a 5 fs integration time step to freely sample the conformational space. The first 20% of the trajectory was discarded as equilibration, and 200 conformations were selected from the remaining trajectory as independent starting structures. For each initial conformation, crosslinks from the respective dataset were applied and the system was first equilibrated at 300 K for 60 ns. Simulated annealing was then performed by decreasing the temperature from 300 K to 0 K in 25 K increments, with 10 ns of simulation at each temperature. The final structures obtained at 0 K constituted the representative conformational ensemble for subsequent analysis.

