## Supplementary Information for "A Phosphorylation Switch Modulates Configurational Codes in the Oncofetal IGF2BP RNA Binding Paralogs"

**Supplementary Figures 1 – 4**

**Supplementary Tables 1 – 4**

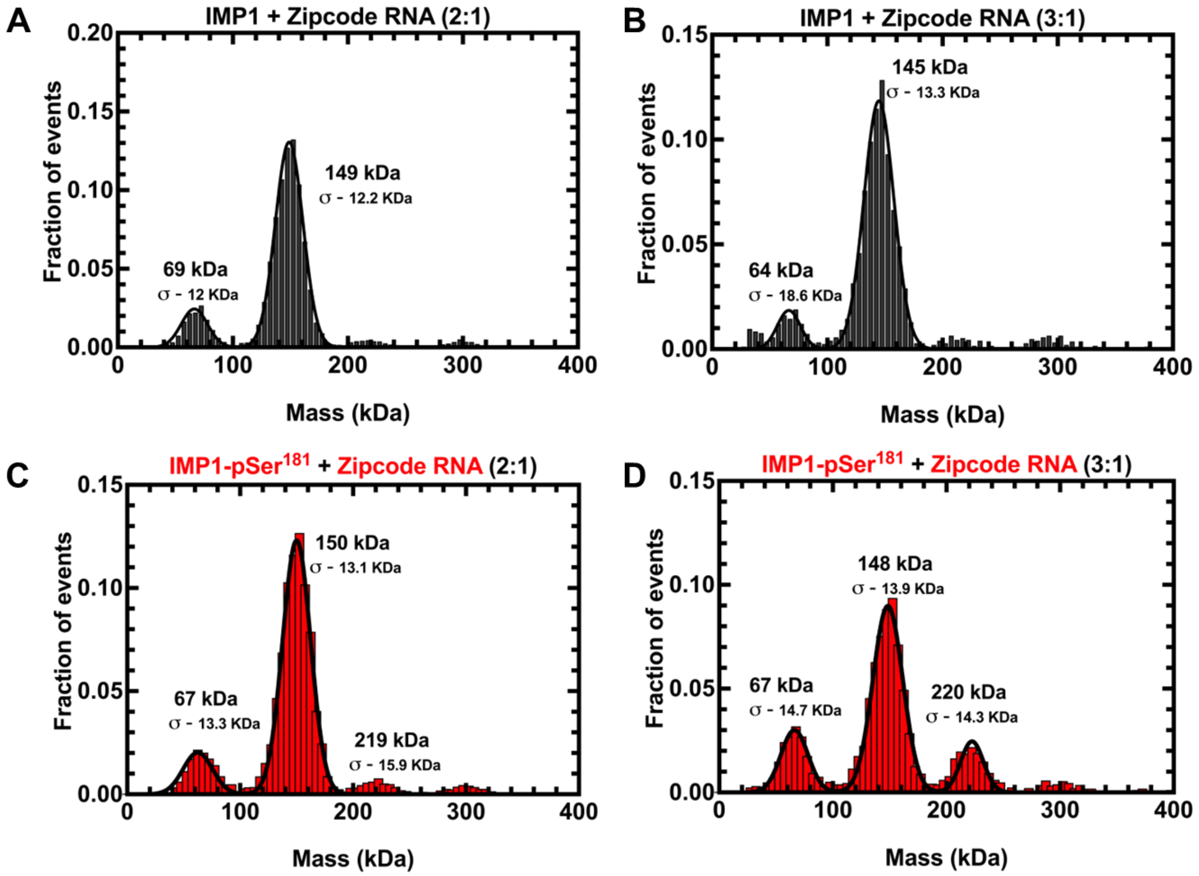


**Supplementary Figure 1. Mass photometry analysis of higher ratios of protein-RNA complexes.** Mass photometry analysis of IMP1 in complex with Zipcode RNA at either **A)** 2:1 or **B)** 3:1 protein:RNA stoichiometry. Mass photometry analysis of IMP1-pSer$^{181}$ in complex with Zipcode RNA at either **C)** 2:1 or **D)** 3:1 protein:RNA stoichiometry. The predicted mass for IMP1 is 64.3 kDa and the Zipcode RNA is 17.6 kDa. The predominant species in all four experiments corresponds to two IMP1 molecules bound to one RNA (~146 kDa). IMP1-pSer-$^{181}$ forms higher order oligomers on RNA compared to IMP1.

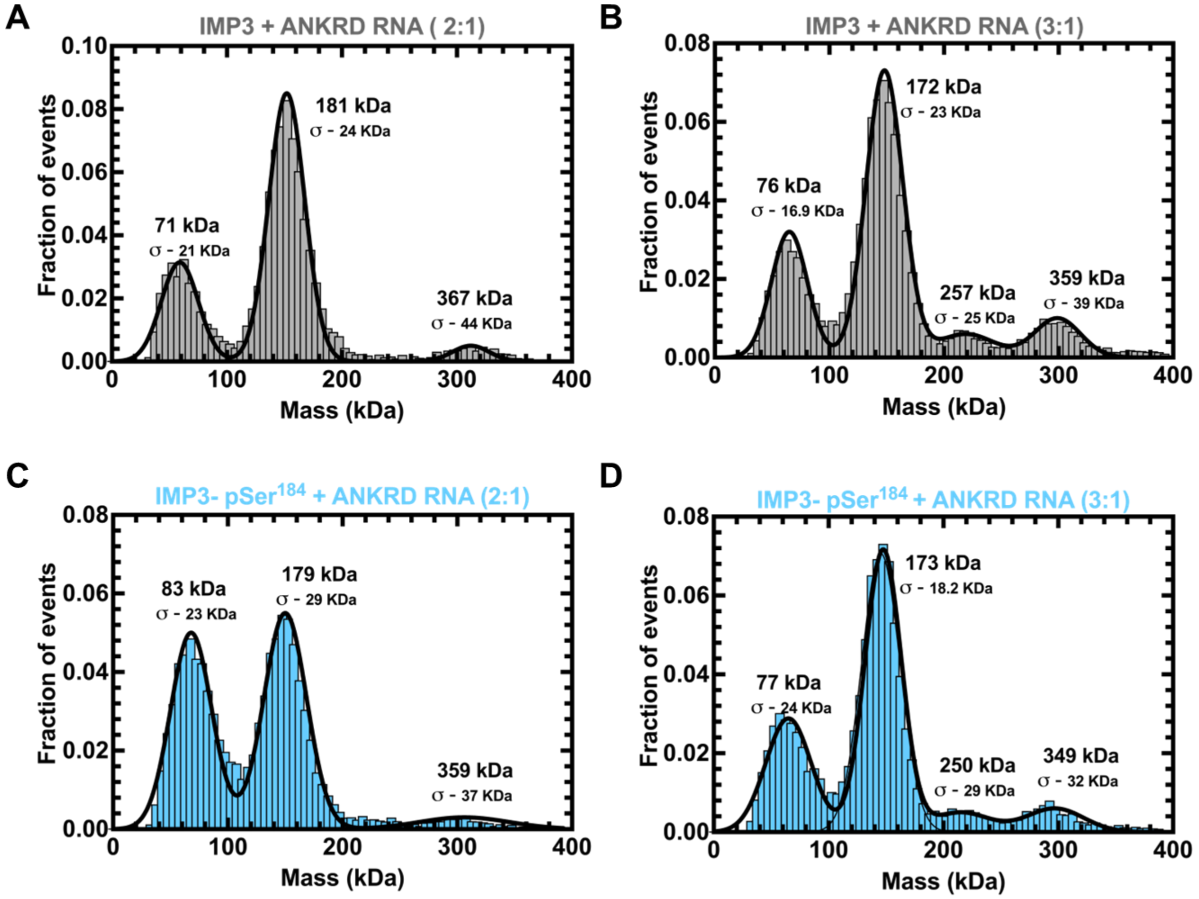


**Supplementary Figure 2. Mass photometry analysis of higher ratios of IMP3-RNA complexes.** Mass photometry analysis of IMP3 in complex with ANKRD RNA at either **A)** 2:1 or **B)** 3:1 protein:RNA stoichiometry. Mass photometry analysis of IMP3-pSer$^{184}$ in complex with ANKRD RNA at either **C)** 2:1 or **D)** 3:1 protein:RNA stoichiometry. The predicted mass for IMP3 is 64.5 kDa and the ANKRD RNA is 38.7 kDa. The first major species in all four experiments appears to be a mixture of free IMP3 (~64.5 kDa) or one IMP3 bound to one RNA molecule (~103 kDa). The second major species is likely two IMP3 molecules bound to one RNA (~167.7 kDa).

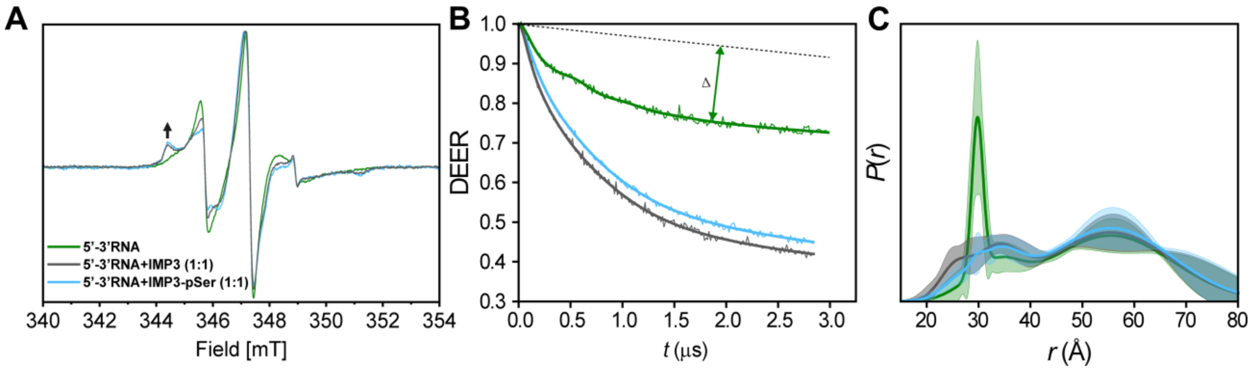


**Supplementary Figure 3. CW EPR and DEER spectroscopy of IMP3 and IMP3-pSer$^{184}$ bound to the non-cognate Zipcode RNA. A)** CW EPR spectra of doubly-labeled RNA in the absence (green) and presence of IMP3 (black), or and IMP3-pSer$^{184}$ (cyan). **B** and **C)** Raw DEER decays and fits are presented for the experimentally determined distance distributions *P(r)*. Confidence bands (2σ) are shown about the best fit lines.

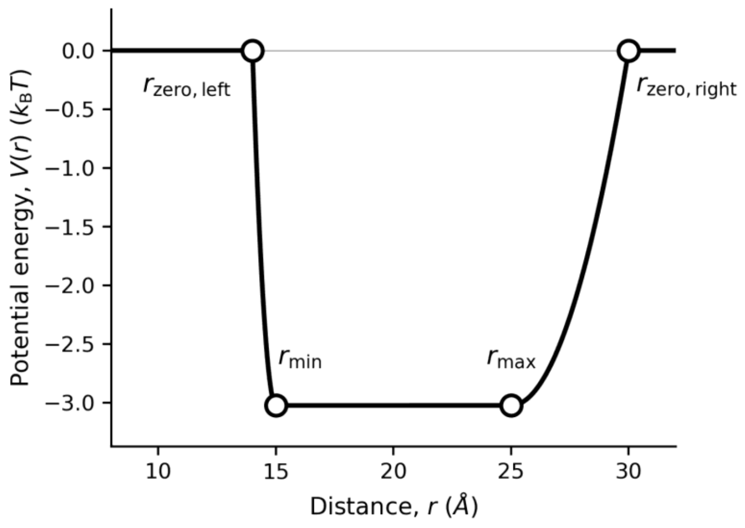


**Supplementary Figure 4. Plot of the potential used to model crosslink interactions.** The potential is defined between a specific i-j pair by an attractive flat-bottom well between ($r_{min}$) and ($r_{max}$) with smooth harmonic transitions at finite-range to zero energy at ($r_{zero,left}$) and ($r_{zero,right}$).

**Supplementary Table 1**

| RNA Oligonucleotide | Sequence |
|---|---|
| Zipcode RNA (IMP1) | 5′ACCGGACUGUUACCAACACCCACACCCCUGUGAUGAAACAAAACCCA<br>UAAAUGC 3′ |
| Spin-labeled Zipcode RNA (IMP1; DEER) | 5'ACC**M**GACUGUUACCAACACCCACACCCCUGUGAUGAAACAAAACCCA<br>UAAA**M**GC3' |
| ANKRD (IMP3) | 5′GGGUCUGUCCGAAGGCAGCUUUUUGUCACAGUUGUGAAGACAUCCAA<br>UCGCCACACAACAACAGUCACAACCACGGCAAGCAACAACAACACUGCA<br>CCCACAAAUGCCACAUAUCCUAUGC 3′ |

**M** denotes the position of the spin label (isothiocyanate derivative of an isoindoline nitroxide) on the RNA. Structure of the spin label is shown below.

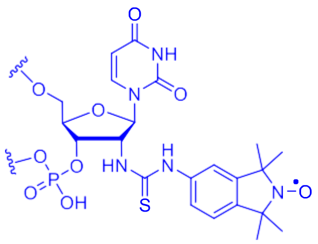

**Supplementary Table 2. Composition of media and reagents used for Phosphoserine incorporation media.**

**A) ZY-non inducing media (ZY-NIM)**

| Components | For 50 mL |
|---|---|
| ZY media | 47.25 mL |
| 1 M $MgSO_4$ | 0.1 mL |
| 25x M-salts | 2 mL |
| 40 % (w/v) α-D-glucose | 0.625 mL |
| Trace metals solution (5000x) | 0.01 mL |

**B) ZY-auto inducing media (ZY-AIM)**

| Components | For 1 L |
|---|---|
| ZY media | 940 mL |
| 1 M $MgSO_4$ | 2 mL |
| 25x M-salts | 40 mL |
| 50x 5052 solution | 20 mL |
| Trace metals solution (5000x) | 0.2 mL |

**C) Composition of 25X M-salts**

| Components | For 1 L |
|---|---|
| Sodium phosphate dibasic | 88.73 g |
| Potassium phosphate dibasic | 85.05 g |
| Ammonium chloride | 66.86 g |
| Sodium sulfate anhydrous | 17.75 g |

**D) 50x 5052 solution**

| Components | For 1 L |
|---|---|
| α-D-glucose | 2.5 g |
| Lactose | 50 g |
| Glycerol (v/v) | 125 mL |

**Supplementary Table 3. Secondary structure composition analysis from circular dichroism.**

| Summary | IMP1 | IMP1-pSer$^{181}$ | IMP3 | IMP3-pSer$^{184}$ |
|---|---|---|---|---|
| Helix | 45.3 ± 1.819 | 51.5 ± 0.306 | 52 ± 0.058 | 45.5 ± 0.200 |
| Antiparallel | 8.7 ± 0.153 | 8.5 ± 0.058 | 6.8 ± 0.115 | 9.1 ± 0.058 |
| Parallel | 0 | 0 | 0 | 0 |
| Turn | 12.2 ± 0.551 | 10.6 ± 0.05 | 10.3 ± 0.132 | 12.2 ± 0.043 |
| Others | 33.8 ± 2.194 | 29.5 ± 0.231 | 30.8 ± 0.021 | 33.2 ± 0.35 |

**Supplementary Table 4. Mass calculations from mass photometry experiments.**

| Protein | No RNA | (Protein: RNA) 1:1 | (Protein: RNA) 2:1 | (Protein: RNA) 3:1 |
|---|---|---|---|---|
| **IMP1** | Peak 1: 57kDa ± 9.5kDa (83%)<br><br>Peak 2: 121kDa ± 19.2kDa, (14%) | Peak 1: 69kDa ± 12.9kDa (12%)<br><br>Peak 2: 152kDa± 12.9kDa (84%) | Peak 1: 69kDa ± 12kDa (14%)<br><br>Peak 2: 149kDa± 12.2kDa (79%)<br><br>Peak 3: 217kDa± 17.6kDa (4%)<br><br>Peak 4: 296kDa± 13.6kDa (3%) | Peak1 : 64kDa ± 18.6kDa (13%)<br><br>Peak 2: 145kDa ± 13.3kDa (77%)<br><br>Peak 3: 214kDa ± 13.6kDa (4%)<br><br>Peak 4: 291kDa ± 14.3kDa (14%) |
| **IMP1-pSer[181]** | Peak1: 63kDa ± 10.7kDa (70%)<br><br>Peak 2: 138kDa ± 12.7kDa (19%)<br><br>Peak 3: 214kDa ± 13.7kDa (5%)<br><br>Peak 4: 299kDa ± 108kDa (6%) | Peak1: 72kDa ± 17.2kDa (12%)<br><br>Peak 2: 153kDa ± 13.8kDa (82%)<br><br>Peak 3: 230kDa ± 22kDa (3%)<br><br>Peak 4: 308kDa ± 16.8kDa (3%) | Peak1: 67kDa ± 13.3kDa (13%)<br><br>Peak 2: 150kDa ± 13.1kDa (77%)<br><br>Peak 3: 219kDa ± 15.9kDa (5%)<br><br>Peak 4: 303kDa ± 76kDa (5%) | Peak1: 67kDa ± 14.7kDa (19%)<br><br>Peak 2: 148kDa ± 13.9kDa (59%)<br><br>Peak 3: 220kDa ± 14.3kDa (15%)<br><br>Peak 4: 298kDa ± 16.2kDa (5%) |
| **IMP3** | Peak1: 70kDa ± 14.9kDa (100%) | Peak1: 65kDa ± 17.1kDa (12%)<br><br>Peak 2: 184kDa ± 30kDa (82%)<br><br>Peak 3: 393kDa ± 49kDa (3%) | Peak1: 71kDa ± 21kDa (26%)<br><br>Peak 2: 181kDa ± 24kDa (66%)<br><br>Peak 3: 367kDa ± 44kDa (7%) | Peak1: 76kDa ± 16.9kDa (20%)<br><br>Peak 2: 172kDa ± 23kDa (55%)<br><br>Peak 3: 257kDa ± 25kDa (6%)<br><br>Peak 4: 359kDa ± 39kDa (10%) |
| **IMP3-pSer[184]** | Peak1: 71kDa ± 11kDa (91%)<br><br>Peak 2: 122kDa ± 31kDa (8%) | Peak1: 72kDa ± 23kDa (41%)<br><br>Peak 2: 175kDa ± 27kDa (53%)<br><br>Peak 3: 345kDa ± 142kDa (6%) | Peak1: 83kDa ± 23kDa (45%)<br><br>Peak 2: 179kDa ± 29kDa (50%)<br><br>Peak 3: 359kDa ± 37kDa (3%) | Peak1: 77kDa ± 24kDa (28%)<br><br>Peak 2: 173kDa ± 18.2kDa (54%)<br><br>Peak 3: 250kDa ± 29kDa (6%)<br>Peak 4: 349kDa ± 32kDa (7%) |
